# Directed evolution expands CRISPR-Cas12a genome editing capacity

**DOI:** 10.1101/2025.03.26.645588

**Authors:** Enbo Ma, Kai Chen, Honglue Shi, Kevin M. Wasko, Isabel Esain-Garcia, Marena I. Trinidad, Kaihong Zhou, Jinjuan Ye, Jennifer A. Doudna

## Abstract

CRISPR-Cas12a enzymes are versatile RNA-guided genome-editing tools with applications encompassing viral diagnosis, agriculture and human therapeutics. However, their dependence on a 5’-TTTV-3’ protospacer-adjacent motif (PAM) next to DNA target sequences restricts Cas12a’s gene targeting capability to only ∼1% of a typical genome. To mitigate this constraint, we used a bacterial-based directed evolution assay combined with rational engineering to identify variants of *Lachnospiraceae bacterium* Cas12a (LbCas12a) with expanded PAM recognition. The resulting Cas12a variants use a range of non-canonical PAMs while retaining recognition of the canonical 5’-TTTV-3’ PAM. In particular, biochemical and cell-based assays show that the variant Flex-Cas12a utilizes 5’-NYHV-3’ PAMs that expand DNA recognition sites to ∼25% of the human genome. With enhanced targeting versatility, Flex-Cas12a unlocks access to previously inaccessible genomic loci, providing new opportunities for both therapeutic and agricultural genome engineering.

## INTRODUCTION

CRISPR-Cas12a is an RNA-guided endonuclease used for genome editing in plants, animals and human cells (1–6). It employs self-processed CRISPR-RNAs (crRNAs) to locate target DNA sequences in the genome (7, 8). Target recognition by Cas12a relies on a 5’-TTTV-3’ (V = A, C or G) protospacer adjacent motif (PAM) as a key signal that enables crRNA-mediated DNA binding and formation of an RNA-DNA R-loop spanning a 20-base-pair (20-bp) target sequence. R-loop formation triggers Cas12a-mediated target DNA cleavage, inducing site-specific sequence changes during the DNA repair process (7, 9, 10).

Although Cas12a is less widely used than Cas9 (11, 12), its unique properties— including its low temperature-tolerant ability to generate staggered double-strand breaks and autonomously process crRNA arrays—make it particularly attractive for applications in plants and for multiplexed genome targeting (7, 8, 13). Furthermore, engineered Cas12a enzymes with improved nuclease activity have expanded their utility across diverse cell types and organisms (14–17). However, Cas12a’s stringent requirement for a four-nucleotide 5’-TTTV-3’ PAM limits targetable sites to ∼1% of a typical genome (18). In comparison, the canonical Cas9 from *Streptococcus pyogenes* (SpyCas9), which requires only a 5’-NGG-3’ PAM, can access >6% of the genome (18). The stringent PAM dependency stems from CRISPR’s natural immune function in bacteria, where PAM recognition helps distinguish foreign DNA from the host genome, ensuring selective cleavage of invasive genetic fragments. Efforts to relax PAM requirements through structure-guided engineering of Cas9 and Cas12a enzymes have generated variants with expanded targeting capabilities (18–21). However, these modifications often reduce cleavage kinetics or increase off-target activity (22, 23), likely due to slower and more promiscuous target search mechanisms (24–26).

To address these limitations, we explored whether a comprehensive mutational approach could lead to Cas12a variants with relaxed PAM requirements and without compromised genome editing efficiency. By combining directed evolution and rational engineering, we generated *Lachnospiraceae bacterium* Cas12a (LbCas12a) variants with both expanded PAM tolerance and robust nuclease activity. Among these, we identified Flex-Cas12a, a standout variant carrying six mutations (G146R, R182V, D535G, S551F, D665N, and E795Q). Flex-Cas12a not only retains efficient cleavage activity at canonical 5’-TTTV-3’ sites but also recognizes a range of PAM sequences (5’-NYHV-3’), thereby expanding potential genome accessibility from ∼1% to over 25%. These improvements provide new tools for genome engineering across diverse biological systems, advancing both fundamental and translational applications.

## MATERIALS AND METHODS

### Library generation of LbCas12a variants with random mutations in the PAM-interacting (PI) and Wedge (WED) domains

For directed evolution (Figure 1A), we chose the region containing both PI and WED domains from *Lachnospiraceae bacterium* Cas12a (LbCas12a) to generate libraries using random mutagenesis (Figure 1B), since these two domains interact with the PAM and its surrounding nucleotides (9, 10, 27, 28).

**Figure 1.**
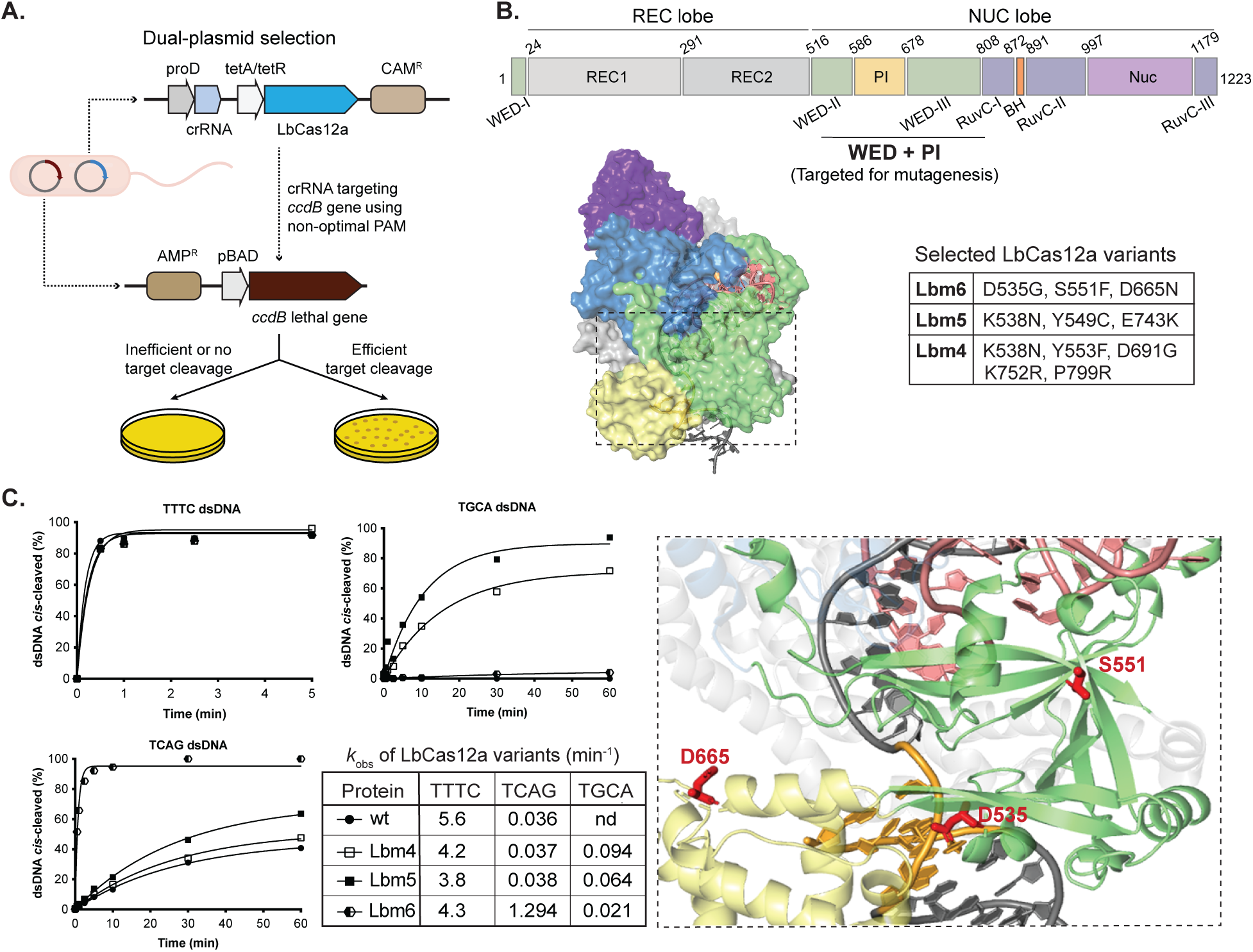
Generation of PAM-relaxed Cas12a enzymatic variants. **A.** Schematic presentation of the dual-plasmid selection system used for directed evolution. Each of four crRNAs (crRNA1-4) was designed for targeting sequences flanked by a randomly designed non-canonical PAMs of AGCT, AGTC, TGCA or TCAG. All DNA target and crRNA sequences are listed in Table S1. **B.** Schematic presentation of the LbCas12a functional domains. The highlighted middle region (WED+PI, top panel) is the target region for mutagenesis. Overall structure (PDB: 5XUS) of LbCas12a-dsDNA-crRNA ternary complex (52) is shown on the left side of the middle panel, while mutations in three selected variants are listed in the table on right side of the middle panel. The bottom panel displays the structural positions of the mutations of D535G, S551F and D665N in Lbm6 and the three mutated residues are labeled in red. The PAM sequence is shown in brown in the structure. **C.** *In vitro* kinetic studies of *cis*-cleavage activity on different PAM DNAs. Cleavage efficiencies of wild-type LbCas12a (wt) and three selected variants (Lbm4, Lbm5, and Lbm6) are shown. Each data point represents the average of two independent experiments. DNA substrates used in these assays are DNA T0 (listed in Table S1) with different PAM sequences. TTTC is a canonical PAM, while others, non-canonical PAMs. Each PAM sequence is listed on the top of each panel. The observed rate constants (*k*_obs_) from these *cis*-cleavage assays are also listed.

To generate a library of random mutations in this target region, we performed error-prone PCR to amplify the DNA fragment encoding PI and WED-II/III domains (Figure 1B) as described elsewhere (16, 27). To limit the error rate to 6- to 9-nucleotide mutations per kilobase, 2.4 µL of 10 mM MnCl_2_ was added to a 100 µL PCR reaction (total volume), which contained 10 µL of 10X ThermoPol reaction buffer, 2 µL of 10 mM primers, 30 ng of template plasmid and 1 µL of ThermoTaq DNA Polymerase (M0267S, NEB). The resulting error-prone PCR products were then used to replace the corresponding wild-type fragment in a LbCas12a bacterial expression plasmid using standard cloning procedures.

### Selection of PAM-relaxed LbCas12a variants using directed evolution

To isolate LbCas12a variants with relaxed PAM requirement, we carried out directed evolution using a dual-bacterial selection system described previously (Figure 1A) (15, 16, 22, 29, 30). Briefly, we constructed four independent chloramphenicol-resistant (CAM^+^) bacterial expression libraries, each harboring the mutagenized PI and WED-II/III domains along with a specific CRISPR RNA (crRNA). Each crRNA was designed to direct cleavage at a target sequence adjacent to a non-canonical PAM (crRNA1: AGCT; crRNA2: AGTC; crRNA3: TGCA; crRNA4: TCAG). These PAMs and target sites were randomly chosen within the *ccdB* lethal gene located in the selection plasmid which contains an Ampicillin-resistant (Amp^+^) gene (Figure 1A; sequences listed in Supplementary Figure S1A and Table S1). For each library, two nanograms of plasmid DNA were electroporated into 50 µl of *E. coli* strain *BW25141*(*DE3*) competent cells containing an ampicillin-resistant (Amp^+^) plasmid encoding an arabinose-inducible *ccdB* lethal gene. The expression of this *ccdB* lethal gene is tightly controlled by the arabinose-inducible promoter pBAD for positive selection (Figure 1A).

After recovery of electroporated bacteria in 2 mL of SOB for 40 min at 37°C, 5 µL of the bacterial culture were plated onto a CAM-containing Petri agar-dish (as a control) and the remaining culture was plated on another Petri agar-dish containing both arabinose and CAM. Positive colonies that grew on plates containing arabinose and CAM were collected and re-plated. Plasmids from individual re-plated colonies were then isolated and sequenced. In this study, two rounds of selection were performed to enrich for LbCas12a variants with relaxed PAM specificity.

### Protein expression and purification

The expression and purification of LbCas12a proteins was carried out using the CL7/IM7Expression and Purification protocol (Trialtus Bioscience) with some modifications. Briefly, the coding sequences for each LbCas12 variant were cloned into a dual-tagged (His/CL7) expression vector and transformed into *E. coli* BL21(DE3) cells (NEB). Cells were cultured in 2xYT medium (Thermo Fisher Scientific) supplemented with 100 µg/mL of ampicillin at 37°C until its optical density (OD_600_) reached 0.6-0.8 (using an overnight starter culture at a 1:40 ratio). The cultures were then cooled on ice for 30 to 60 min before induction with 0.5 mM isopropyl β-D-1-thiogalactoside (IPTG). Following IPTG induction, cells were grown overnight at 16°C. To purify the Cas12a proteins, the cultured cells were harvested and resuspended in lysis buffer (50 mM Tris-HCl (pH 7.5), 20 mM imidazole, 1M NaCl, 10% (v/v) glycerol and supplemented with 1 mM tris(2-carboxyethyl)phosphine (TCEP) and a cOmplete^TM^ Protease Inhibitor Cocktail Tablet (Millipore Sigma) for every 50 ml). Lysis was conducted by sonication. The lysed cultures were clarified by centrifugation for 60 min at 18,000 xg. The clarified supernatant was then applied to Ni-NTA resin (Qiagen), which was pre-equilibrated with wash buffer (50 mM Tris-HCl (pH 7.5), 20 mM imidazole, 0.5 mM TCEP, 1 M NaCl). The mixture was then incubated for 30-60 min at 4°C to allow binding of the His-tagged target protein to Ni-NTA resin, then washed three times with the same buffer, and finally eluted with a buffer containing 50 mM Tris-HCl (pH 7.5), 300 mM imidazole, 1 M NaCl, 10% glycerol and 1 mM TCEP. The Ni-NTA resin-purified proteins were applied 2-3 times to Im7 beads. The beads were then washed with 5-10 column volumes of a wash buffer containing 20 mM HEPES (pH 7.5), 1 M NaCl, 10% glycerol and 1 mM TCEP before addition of Pierce human rhinovirus (HRV) 3C protease (Thermo Fisher Scientific) to release the LbCas12a protein. The solution containing target protein was collected and then concentrated with a 30,000 MWCO concentrator (Millipore Sigma). The concentrated protein was further purified by application to a Superdex200 Increase 10/300 GL (Cytiva) column using gel filtration buffer (20 mM HEPES (pH 7.5), 150 mM KCl, 10% glycerol and 1 mM TCEP). Peak fractions containing Cas12a proteins were collected, concentrated and quantified using a NanoDrop 8000 Spectrophotometer (Thermo Fisher Scientific), and stored at −80°C after flash-freezing with liquid nitrogen.

### Nucleic acid preparation

All DNA and RNA oligos used for *in vitro* experiments in this study were purchased from Integrated DNA Technologies, Inc. (IDT) and HPLC or PAGE-purified. For genome editing experiments, crRNAs contained chemical modifications at their 3’- or 5’-ends to improve stability and editing efficiency in cells (detailed in IDT guidance).

DNA substrate (T0, shown as non-target strand) used for *in vitro* cleavage assays is 5’-GACGACAAAAC**NNNN**GATCGTTACGCTAACTATGAGGGCTGTCTGTGGAATGCTA-3’ (here, NNNN is a PAM sequence and listed in each corresponding figure). Guide RNA of crRNA0 used for *in vitro* cleavage assays is 5’-AAUUUCUACUCUUGUAGAUGAUCGUUACGCUAACUAUGAGGGC-3’.

Four DNA targets used in the directed evolution selection are listed below (bold letters indicate non-canonical PAM sequences in front of the protospacers which are underlined):

T1: **AGCT**TTCATCCCCGATATGCACCACCGG,

T2: **AGTC**TCCCGTGAACTTTACCCGGTGGTG,

T3: **TGCA**TATCGGGGATGAAAGCTGGCGCAT, and

T4: **TCAG**ATAAAGTCTCCCGTGAACTTTACC.

Their corresponding guide RNAs (crRNAs) used for the selection are crRNA1, crRNA2, crRNA3 and crRNA4, respectively.

The sequences of all DNAs and RNAs used for *in vitro* assays in this study are also listed in Supplementary Table S1.

### DNA cleavage assays

Typical *cis*-cleavage assays (otherwise will be stated) were carried out in a buffer containing 60 nM protein, 72 nM of crRNA and 10 nM of 5’-FAM-labeled non-target-strand of a target dsDNA in cleavage buffer (20 mM HEPES (pH 7.5), 150 mM KCl, 10 mM MgCl_2_, 1% glycerol and 1 mM TCEP). Specifically, the protein and a guide crRNA were first incubated for 15 min at room temperature to form RNPs and then incubated at 37°C for certain duration of time after addition of labeled target DNA. For *trans*-cleavage assays, the reaction conditions were the same as *cis*-cleavage reactions except that 45 nM unlabeled target dsDNA instead of labeled target dsDNA were used. After incubation for 30 min at 37°C, a labeled random ssDNA (no homology with the target DNAs or crRNAs) was added to the reactions. The reactions were continually incubated for certain durations of time. The reactions of both *cis*- and *trans*-cleavage were quenched with 1X DNA loading buffer (45% formamide, 15 mM EDTA, trace amount of xylene cyanol and bromophenol blue). After denaturation at 95°C for 3 min, the cleavage products were separated using a 15% urea-PAGE gel and quantified using a Typhoon phosphorimager (Amersham, GE Healthcare). Cleavage kinetics were determined by fitting time-course data to the equation:

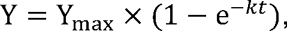

where Y_max_ is the pre-exponential factor, *k* is the rate constant (min^−1^), and t is the reaction time (min).

### PAM depletion assays

A single DNA oligo containing a randomized 6N PAM region (at each position, N = A, C, G, or T) in front of the Cas12a target sequence was synthesized by IDT. The single DNA oligos were annealed and primer-extended into dsDNA oligos using PCR polymerase. To generate a random PAM plasmid library, the dsDNA oligos were subsequently Gibson-cloned into a pUC19 plasmid backbone.

PAM depletion assays were performed under conditions similar to the standard DNA cleavage assays described above. Briefly, 60 nM wild-type LbCas12a or its variants were pre-incubated with 72 nM crRNA complementary to the corresponding protospacer sequence which is downstream of the 6N-PAM region in the plasmid library (protospacer and guide RNA sequences are listed in Supplementary Table S1). The Cas12a-crRNA complex was then added into the plasmid library (13 nM) and incubated at 37°C for 10 minutes. The reaction was quenched by addition of one volume of 2X quench buffer (95% formamide and 30mM EDTA) and heat treatment for 10 min at 95°C. The plasmid DNAs were purified using magnetic beads. After clean-up, the PAM-region in the un-cleaved plasmid DNAs was amplified by PCR in the presence of the primers containing partial Illumina adapter sequences (primer sequences listed in Supplementary Table S1) which allow for subsequent indexing in second-round PCR. The first-round PCR reactions were treated with addition of 0.5 µL of DpnI for 30 min at 37°C before they were purified using magnetic beads. The purified PCR products were subjected to the indexed PCR to make indexed libraries. The indexed libraries were pooled equimolarly, quantified by Qubit and sequenced on an Illumina NextSeq platform (P1 flow cell) with 2×150 bp paired-end reads (IGI NGS sequencing core, UC Berkeley). The sequencing run typically achieved sufficient coverage (> 500,000 reads per library) to provide a comprehensive view of PAM preferences across the 6N library.

Sequencing data were processed according to the high-throughput PAM determination assay (HT-PAMDA) pipeline (31) using a customized Python script provided in (https://github.com/kleinstiverlab/HT-PAMDA). In brief, raw FASTQ reads were aligned to a reference sequence to extract the 6-nt PAM regions. For each PAM, its frequency in the Cas12a RNP-treated sample was compared to that in a control sample (plasmid without treatment with any RNP) to calculate a relative depletion rate after a single 10-min reaction. These depletion rate constants (expressed on a log scale) serve as a quantitative measure of cleavage activity at each PAM site. The resulting heatmaps display the log rate constants using a color gradient, where darker hues indicate higher cleavage activity (or greater depletion) and lighter hues denote lower activity.

### Cell culture

HEK293T cells (UC Berkeley Cell Culture Facility) were cultured using Dulbecco’s Modification of Eagle’s Medium (DMEM) with L-glutamine, 4.5 g/L glucose and sodium pyruvate (Corning) plus 10% FBS, and penicillin and streptomycin (Gibco).

Neural progenitor cells (NPCs) were isolated from embryonic day 13.5 Ai9-tdTomato homozygous mouse brains. Cells were cultured as neurospheres in NPC medium (DMEM/F12 with glutamine, Na-pyruvate, 10 mM HEPES, non-essential amino acids, 1x penicillin and streptomycin, 1x 2-mercaptoethanol, B-27 without vitamin A, N2 supplement, and growth factors, bFGF and EGF (both 20 ng/ml at final concentration)). NPCs were passaged using MACS Neural Dissociation Kit (Papain, CAT# 130-092-628) following the manufacturer’s protocol. bFGF and EGF were refreshed every three days and cells were passaged every six days. The NPCs were authenticated by immunocytochemistry marker staining for Nestin and GFAP.

### Genome editing

To evaluate the genome editing activity of LbCas12a variants, we electroporated the corresponding ribonucleoproteins (RNPs) into regular HEK293T cells, EGFP-expressing HEK293T cells (HEK293T-EGFP), or tdTomato-transgene-containing neural progenitor cells (NPCs). For each transfection, 100 pmol of LbCas12a protein was pre-incubated with 120 pmol crRNA at room temperature for 15-25 min to form RNPs, after which 80 pmol of a ssDNA electroporation enhancer (purchased from IDT) was added.

For HEK293T or HEK293T-EGFP cells, RNP nucleofection was carried out with Lonza (Allendale, NJ) SF cell kits in an Amaxa 96-well Shuttle system (program code CM-130). Each nucleofection reaction consisted of approximately 2.0×10^5^ cells and 100 pmol RNP in a total volume of 20 µL of supplemented SF buffer following the manufacturer’s instructions. After nucleofection, 120 µL of growth medium was added to each nucleofection cuvette, and 5,000–10,000 cells were subsequently transferred to individual wells of a 96-well tissue culture plate preloaded with 100–120 µL of full medium. After incubation at 37°C for 3 days, the cell culture medium was refreshed. After an additional 3 days of culture, flow cytometry was used to determine editing efficiency, as indicated by loss of either EGFP or B2M.

Briefly, HEK293T cells were detached by addition of 30 µL 0.05% Trypsin-EDTA per well and incubated for ∼3 min before addition of 120 µL complete media to each well. 120 µL of the treated cells were transferred to a 96-well U-bottom plate, to which 80 µL of PBS was added in each well and centrifuged at 1,600 rpm for 5 min. After the media were decanted, the cells were washed with 150 µL PBS/BSA (1% BSA). The washed cells were resuspended with 50 µL of staining master mix containing anti-B2M-APC (Biolegend, Luxembourg, 1:200 dilution with PBS supplemented with 1% BSA), or resuspended in PBS to be analyzed directly, when assessing EGFP knockout. The cells were stained for 30 min in the dark before the addition of 100 µL PBS/BSA for washing. Following an additional wash with 100 µL PBS/BSA, cells were resuspended in 150 µL PBS and analyzed by flow cytometry using an Attune Flow Cytometer (Thermo Fisher Scientific).

Nucleofection of NPCs with RNPs was performed using Lonza (Allendale, NJ) P3 cell kits in an Amaxa 96-well Shuttle system with program code EH-100. Each nucleofection reaction consisted of approximately 2.5×10^5^ cells and 100 pmol RNP (preassembled in a phosphate buffer (25 mM NaPi, 300 mM NaCl, 200 mM trehalose, pH 7.5) with a total volume of 20 µL in the supplemented P3 buffer following the manufacturer’s instructions. After nucleofection, 80 µL of growth medium was added to the nucleofection cuvette and 5 µL of NPC culture was then transferred to 96-well tissue culture plates pre-coated (using laminin, fibronectin, and poly-DL-ornithine), with a total culture volume of 100 µL/well. Following a 3-day incubation at 37°C, the medium was refreshed, and the cells were harvested after an additional 3 days for FACS of tdTomato expression.

### Base-editing

We constructed both cytosine base editor (CBE) and adenine base editor (ABE) versions of LbCas12a by modifying two existing plasmid backbones. The CBE backbone (15) is derived from pCAG-NLS(SV40)x2-rAPOBEC1-gs-XTEN-gs-hdLbCas12a(D832A)-gs-UGI-NLS(SV40) (LbBE1.4, Addgene plasmid #114086, RTW1293). The ABE backbone (32) is made from pCMV-dCas12a-ABE8e (Addgene plasmid #193645). LbCas12a base editors (BEs) used human codon-optimized sequences of LbCas12a and its variants for human cells through gBlocks synthesized by Twist Bioscience.

Each base editor plasmid was co-transfected with a corresponding crRNA expression plasmid driven by the human U6 promoter. We designed 4 crRNA constructs for cytosine base editing and 4 crRNA constructs for adenine base editing; their target sequences and PAMs are listed in Supplementary Table S2. For each transfection, 100 ng of the base editor plasmid and 40 ng of the crRNA plasmid were introduced into 2.0×10⁴ HEK293T cells plated in 96-well plates. Transfections were carried out using FuGENE transfection reagent according to the manufacturer’s recommended protocol (Promega). On Day 4 post-transfection, genomic DNAs were isolated according to protocols described below.

### NGS sequencing

To produce amplicons used in NGS, all genomic DNAs were extracted following the protocol supplied in Quick Extraction solution (Epicentre, Madison, WI). Briefly, after the culture medium was removed, 100 µL of Quick Extraction solution were added to each well to lyse the cells (65°C for 20 min and then 95°C for 20 min). The concentration of genomic DNA was determined by NanoDrop and was stored at −20°C.

DNA sequences of potential off-targets were identified as described elsewhere (27). Briefly, we used Cas-OFFinder (v3.0.0b3) (33) and the human reference genome GRCh38.p14 to identify potential off-target sequences for two target sites with canonical PAM of TTTA or the non-canonical PAM of TCAG, respectively. The parameters used in this study to select potential off-target sites are the sequences with ≤1 bulge, ≤5 mismatches and specific PAMs.

To confirm genome editing based on B2M knockout, to investigate off-target editing by the LbCas12a variants in HEK293T cells and to analyze the efficiency of base editing, target or off-target amplicons were PCR-amplified in the presence of corresponding primers. Target loci were first amplified by a first PCR step using locus-specific primers (primer sequences listed in Supplementary Tables S2-S3), and indexed libraries were generated through a second PCR step to append Illumina adapter sequences and unique sample barcodes. The PCR products were purified with magnetic beads (Berkeley Sequencing Core Facility) and then were pooled at equimolar concentrations, quantified, and sequenced on an Illumina NextSeq platform (P1 flow cell) with 2×150 bp paired-end reads at the IGI NGS sequence core, resulting in sufficient coverage of the amplicon (> 10,000 reads per library).

Data were analyzed with Geneious Prime (v2024.07) (https://www.geneious.com/) and CRISPResso2 (v2.2.6) (34) for indel quantification, with a 50% minimum alignment score and two biological replicates for indel analysis and three biological replicates for BE analysis. Paired-end Illumina sequencing reads were initially processed in Geneious Prime (https://www.geneious.com/prime) using the BBDuk module to remove low-quality bases (minimum score of 20) and discard reads shorter than 20 nucleotides. The filtered reads were then merged with BBmerge, also within Geneious Prime, to produce contiguous sequences where possible. For samples that included only forward (R1) reads, no merging step was performed. After preprocessing, the resulting reads were analyzed in CRISPResso2 (https://github.com/pinellolab/CRISPResso2) to quantify both indel formation and base editing frequencies. Two commands were used, one focusing on indel rates and another on base editing outcomes:

*CRISPResso --fastq_r1 MERGED_READS --amplicon_seq AMPLICON_SEQUENCE -- guide_seq GUIDE_SEQUENCE -n nhej -wc -3 -w 5 --plot_window_size 20 -o OUTPUT_FILE*

*CRISPResso --fastq_r1 MERGED_READS --amplicon_seq AMPLICON_SEQUENCE -- guide_seq GUIDE_SEQUENCE -n nhej -wc -12 -w 6 --plot_window_size 20 – base_editor_output -o OUTPUT_FILE*

In these commands, *--fastq_r1* points to the processed FASTQ file (merged or unmerged), while *--amplicon_seq* indicates the entire amplified region being examined. The *-- guide_seq* parameter specifies the guide RNA spacer. The flags *-wc* and *-w* set the center and size of the quantification window relative to the 3’ end of the guide RNA, and *-- plot_window_size* affects the visualization range for indels. When base editing was examined, *--base_editor_output* was included, allowing CRISPResso2 to report C-to-T or A-to-G substitutions over the defined window. This workflow provided a clear assessment of both indel formation and base editing efficiency for the targeted loci.

## RESULTS

### Directed evolution of LbCas12a variants with relaxed PAM requirements

The requirement for T-rich PAMs (5’-TTTV-3’, where V is A, C or G) in LbCas12a limits its applications in genome editing (11, 18). To overcome this limitation, we employed a bacterial selection-based directed evolution system (15, 16, 22, 30) to generate LbCas12a variants capable of recognizing a wide spectrum of PAM sequences (Figure 1A). Guided by previous findings that the PAM-interacting (PI) and wedge (WED) domains in both Cas12a and Cas9 are critical for PAM binding and DNA interaction (9, 10, 27), we targeted the PI, WED-II and WED-III regions of LbCas12a for random mutagenesis via error-prone PCR (Figure 1B, top panel).

We arbitrarily designed four guide crRNAs (crRNA1-4) to target DNA sequences flanked by non-canonical PAMs of 5’-AGCT-3’, 5’-AGTC-3’, 5’-TGCA-3’ and 5’-TCAG-3’, respectively (Supplementary Figure S1A). These DNA target sequences are located within the coding sequence of the lethal protein *ccdB* in the selection plasmid (16) (Figure 1A). Following two sequential rounds of selection, seven PAM-relaxed LbCas12a variants (designated Lbm1 to Lbm7) emerged from the libraries corresponding to crRNA2, crRNA3, and crRNA4 (Figure 1B, middle panel, right; Supplementary Figure S1B). Specifically, four LbCas12a variants (Lbm1-4) were selected from crRNA2, one (Lbm5) from crRNA3 and two (Lbm6-7) from crRNA4 (Supplementary Figure S1B). Interestingly, no variant was obtained with guide RNA of crRNA1, likely because the corresponding 5’-AGCT-3’ PAM resembles part of the crRNA scaffold (5’-AGAU-3’), which could lead to self-targeting of the crRNA1 locus. Most of the mutations in the selected LbCas12a variants occurred in the WED-II and -III domains (Supplementary Figure S1B). The finding that mutations in the WED domains are involved in the alteration of LbCas12a PAM-binding capabilities (Figure 1B, middle panel, right; Supplementary Figure S1B) underscores the potential of WED domains to affect target recognition (9, 10, 27, 35). Structural mapping of these mutations onto the known LbCas12a ternary structure showed that while a few mutations were positioned near PAM-recognition sites, others were found in regions previously implicated in crRNA binding or general DNA interactions (Figure 1B; Supplementary Figure S1C). This diverse mutational landscape contrasts with structure-guided rational engineering targeting residues directly involved in PAM binding (18–21, 27, 36). Instead, this random mutagenesis approach aimed to uncover beneficial mutations across multiple protein regions, thereby expanding the potential for engineering Cas12a variants with relaxed PAM requirements.

The seven LbCas12a variant proteins were expressed and purified (Supplementary Figure S1D) to confirm their PAM recognition preferences. Based on biochemical cleavage activities using both short synthetic dsDNA (Supplementary Figure S1E) and plasmid substrates (Supplementary Figure S1F), we chose three variants (Lbm4-6) for detailed kinetic analysis with dsDNA substrates bearing various PAMs. All three variants exhibited cleavage rates comparable to those measured for wild-type LbCas12a (wt) when tested with a canonical 5’-TTTV-3’ PAM containing substrate (*k*_obs_ = 3.8-5.6 min^−1^) and could recognize non-canonical PAMs as well (Figure 1C). Additional cleavage assays revealed that Lbm6 had the most relaxed PAM specificity (Supplementary Figure S1G). Given its expanded target range and robust biochemical activity, we focused on Lbm6 in subsequent studies.

### Construction of a composite genome editor with a relaxed PAM requirement

To assess the genome editing activity of variant Lbm6, we first employed tdTomato reporter neural progenitor cells (NPCs) derived from Ai9 mice (6, 37) and delivered Cas12a ribonucleoproteins (RNPs) via nucleofection (Figure 2A, upper panel). In this system, genome editing at the transgene locus to remove the stop cassette upstream of the tdTomato gene activates tdTomato expression. Six days post nucleofection of RNPs, Lbm6, but not wild-type Lbcas12a, successfully edited loci flanked by non-canonical PAMs (Figure 2A, lower panel). However, at a target site bearing a canonical 5’-TTTA-3’ PAM, Lbm6 generated half the level of editing (34%) as that observed with wild-type LbCas12a (67%) (Figure 2A, lower panel). The lower genome editing rate shown in Lbm6 suggested that mutations in this variant might compromise its activity.

**Figure 2.**
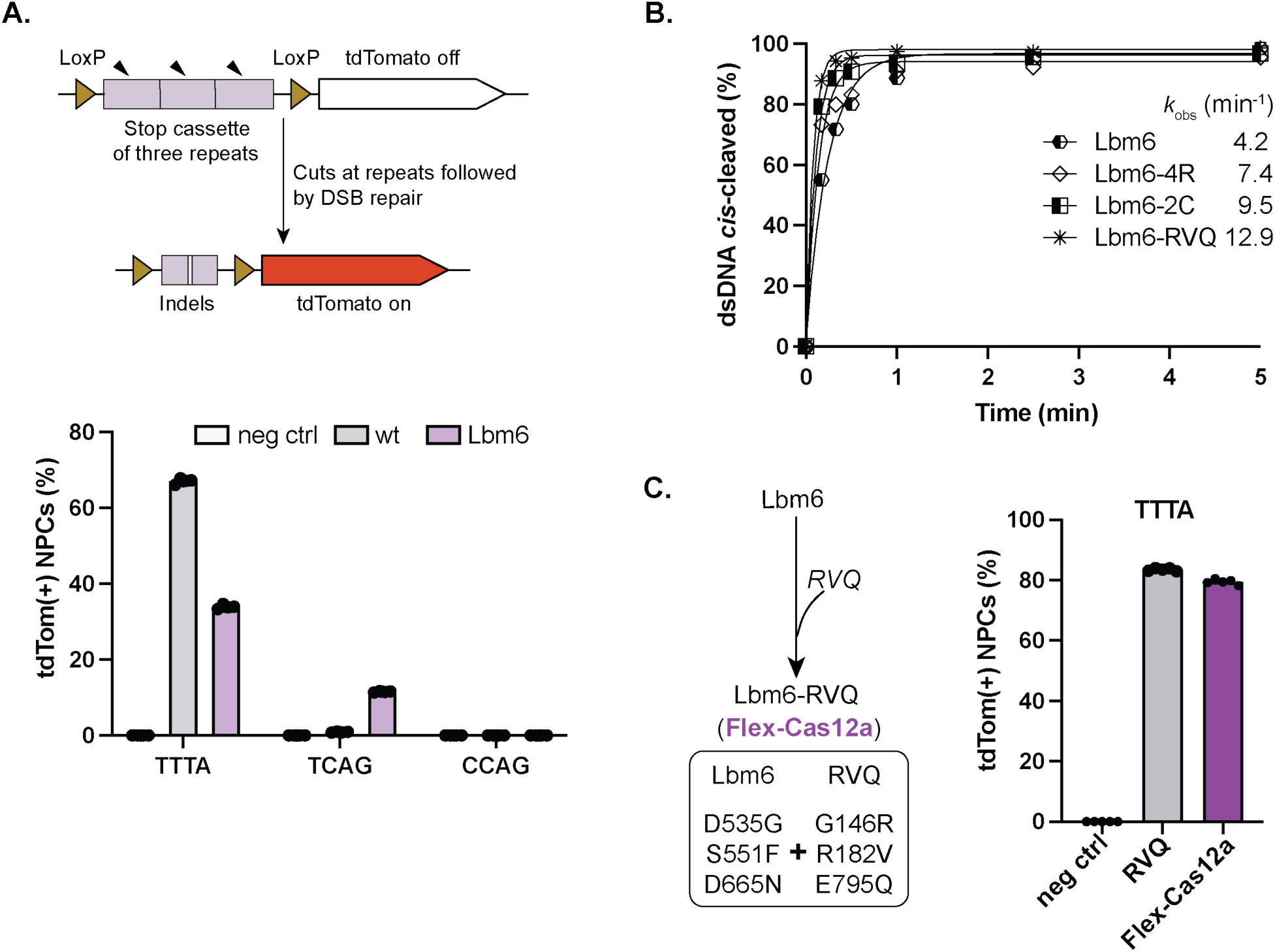
Nuclease activities of Lbm6 and its derivatives. **A.** Genome editing in Ai9 mouse derived neural progenitor cells (NPCs). The upper panel provides a schematic illustration of desired genome editing to turn on the tdTomato transgene in NPCs. The lower panel presents genome editing efficiencies at three target sites in NPCs. PAMs are shown on X-axis. TTTA is a canonical PAM, while others, non-canonical PAMs. Each bar represents the average of four independent experiments. **B.** Improvement of Lbm6 enzymatic cleavage activity. *cis*-cleavage activity of Lbm6 is improved by introduction of the mutations from the activity-enhanced variant of Cas12a-4R, -2C or -RVQ. Introduction of RVQ mutations (G146R, R182V and E795Q) into Lbm6 makes it most active as shown by its *k*_obs_. From now on, the combined version of Lbm6 and RVQ mutations will be named Flex-Cas12a. Each data point represents the average of two independent experiments. DNA target used in this assay is DNA T0 with a canonical PAM of 5’-TTTC-3’. **C.** Genome editing efficiency of Flex-Cas12a. Flex-Cas12a exhibits significantly enhanced genome editing activity which is comparable to LbCas12a-RVQ, at a locus with canonical PAM of 5’-TTTA-3’ in tdTomato NPCs. Negative control (neg ctrl) conditions in **(A** and **C)** means cells which were not treated by any RNP. Each bar represents the average of four independent technical replicates.

Previous studies showed that modifications in either the NUC or REC lobe of Cas12a can significantly enhance its nuclease activity (14, 16, 17). Based on these findings, we tested whether Lbm6’s editing activity could be improved by incorporating known activity-enhancing mutations. Three sets of mutations from Cas12a-RVQ (17), hyperCas12a (14), or iCas12a (16) were introduced into Lbm6, resulting in the new variants Lbm6-RVQ, Lbm6-4R and Lbm6-2C, respectively. Biochemical DNA cleavage assays demonstrated that Lbm6-RVQ displayed the highest activity among the three Lbm6 derivatives (Figure 2B; Supplementary Figure S2A-B). We tested the genome editing activity of this variant in tdTomato NPCs at a locus with a 5’-TTTA-3’ PAM. At this locus, Lbm6-RVQ achieved an 80% editing rate, substantially higher than its parental variant Lbm6 (34%) and wild-type LbCas12a (67%), and comparable to LbCas12a-RVQ (84%) (Figure 2C). Henceforth, we refer to Lbm6-RVQ as Flex-Cas12a in this study.

Given that Cas12a enzymes typically possess both *cis*-cleavage activity and target-activated *trans*-DNA cutting activity (38, 39), we also tested both *cis*- and *trans*-cleavage efficiencies of Flex-Cas12a in comparison with LbCas12a-RVQ. We found that Flex-Cas12a maintained similar *cis*- and *trans*-cleavage activities to LbCas12a-RVQ when tested with canonical 5’-TTTV-3’ PAM targets (Figure 3A, B; Supplementary Figure S3A-B).

**Figure 3.**
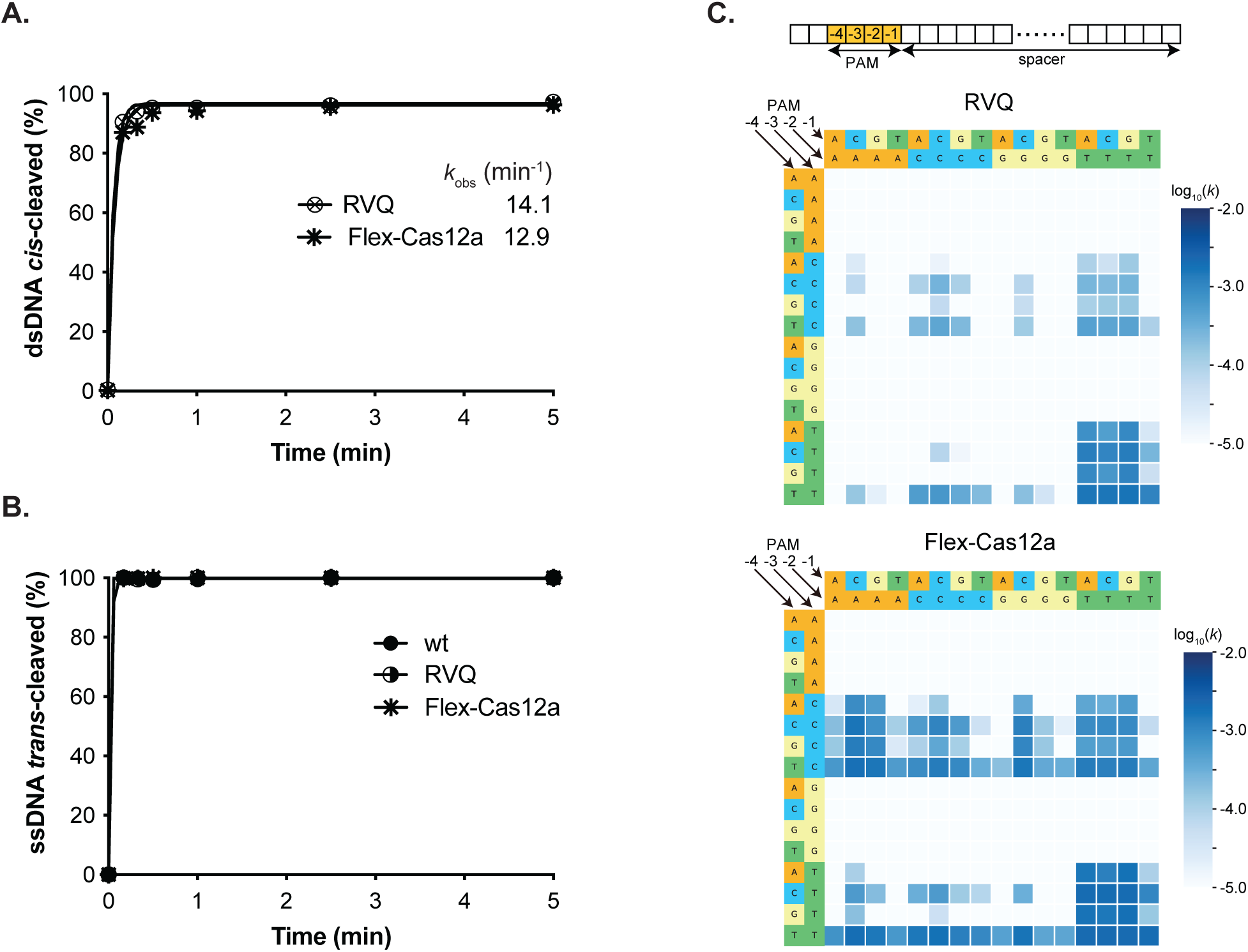
*In vitro* kinetic studies and PAM depletion assays. **A.** *In vitro* kinetic analysis of *cis*-cleavage activity. The observed *k*_obs_ indicate that Flex-Cas12a exhibits cleavage activity comparable to LbCas12a-RVQ (RVQ). Each data point represents the average of two independent experiments. The Target DNA used in this assay is DNA T0 (listed in Table S1) with a canonical PAM of 5’-TTTC-3’. **B.** *In vitro* kinetic analysis of *trans*-cleavage activity. Due to the rapid cleavage rates, the *k*_obs_ of *trans*-cleavage could not be accurately determined from these assays. Each data point represents the average of two independent experiments. Target DNA and guide RNA used in this assay are listed in Table S1. **C.** Expanded PAM recognition by Flex-Cas12a. Heatmaps of NGS results from PAM depletion assays illustrate PAM preferences of LbCas12a-RVQ (RVQ) and Flex-Cas12a. Top panel presents a schematic illustration of nucleotide positions in the PAM motif. Middle panel shows the heatmap for LbCas12a-RVQ, while bottom panel depicts the heatmap for Flex-Cas12a. The heatmap represents the log_10_(k) of the *in vitro* cleavage rate constant (s^−1^).

To systematically analyze the PAM recognition profile of Flex-Cas12a, we performed PAM-depletion assays. While, as previously reported (17), LbCas12a-RVQ primarily recognizes canonical 5’-TTTV-3’ PAMs (Figure 3C, middle panel), Flex-Cas12a utilized a wide range of PAMs, specifically those with sequence 5’-NYHV-3’ (Y is T or C, H is A, C or T, N is any base) (Figure 3C, bottom panel). This expanded recognition enables Flex-Cas12a to potentially target more than 25% of a human genome, extending the ∼6% accessibility achieved at maximum by previous LbCas12a variants (18, 20, 21).

### Specificity of genome editing by Flex-Cas12a in different cell types

To evaluate the utility of Flex-Cas12a at target sites with different PAMs in different mammalian cell types, we conducted genome-editing assays in HEK293T-EGFP cells and tdTomato-expressing mouse neural progenitor cells (NPCs) (16, 27, 40). For these assays, we designed crRNAs targeting ten different reporter genomic loci bearing various non-canonical PAMs in HEK293-EGFP cells and ten such targets in tdTomato NPCs. Six days post-nucleofection of cells with corresponding RNPs prepared with either Flex-Cas12a or LbCas12a-RVQ, flow cytometry analysis showed that Flex-Cas12a edited these targets at levels ranging from 60-80% in both cell types, whereas LbCas12a-RVQ exhibited minimal activity at the loci with non-canonical PAMs (Figure 4A, B; Supplementary Figure S4A, B). These results indicate that Flex-Cas12a can effectively recognize 5’-NYHV-3’ PAMs, significantly expanding its genome editing capability beyond that displayed by wild-type and engineered LbCas12a enzymes (18, 20, 21, 41, 42).

**Figure 4.**
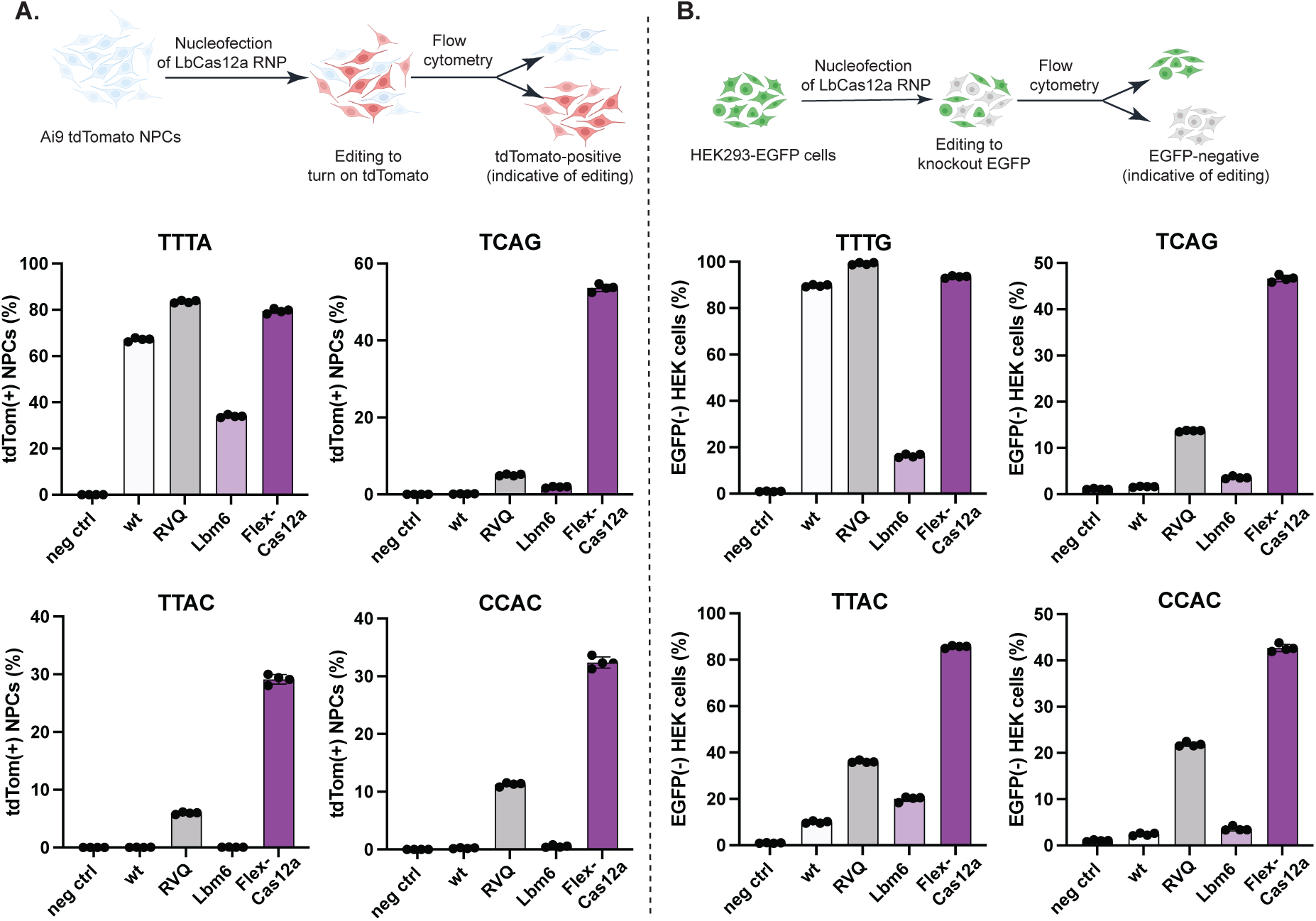
Genome editing in Ai9 tdTomato NPCs and HEK293T-EGFP cells. **A.** Genome editing in tdTomato NPCs. The upper panel provides a schematic illustration of desired genome editing to turn on the tdTomato transgene in NPCs. The lower panels show the genome editing efficiencies at four loci by wild-type (wt), LbCas12a-RVQ (RVQ), Lbm6 and Flex-Cas12a. PAM sequence of each target is shown on top of each panel. **B.** Genome editing in HEK293T-EGFP cells. The upper panel provides a schematic illustration of desired genome editing to turn off the EGFP transgene in HEK293T cells. The lower panels show the genome editing efficiencies of four proteins at four loci. TTTA and TTTG are canonical PAMs, while others, non-canonical PAMs. PAM sequence of each target is shown on top of each panel. The genome editing results from both cell types demonstrate that Flex-Cas12a is significantly more active than its parent version (Lbm6) and can efficiently recognize a broad range of PAM sequences for genome editing. Data are presented as mean ± SD from four independent technical replicates.

Next, we assessed the ability of Flex-Cas12a RNPs to edit endogenous genomic loci by designing crRNAs to target five sites in the human β2-microglobulin (B2M) gene, encoding a broadly expressed surface protein with clinical relevance for cell therapies (43). Knockout was monitored using a B2M-specific monoclonal antibody that indicates successful gene disruption (44). Six days post-nucleofection with Flex-Cas12a RNPs, flow cytometry analysis showed editing efficiencies ranging from 5 to 40% at genomic sites containing either a canonical or non-canonical PAM. In contrast, similar experiments conducted using LbCas12a-RVQ RNPs showed editing only at sites containing canonical 5’-TTTV-3’ PAMs (Figure 5A). Next-generation sequencing (NGS) confirmed that Flex-Cas12a achieved ∼50% editing using a non-canonical TCAG PAM sequence, whereas LbCas12a-RVQ showed minimally detectable activity at this site (Figure 5B). Moreover, off-target editing in both LbCas12a-RVQ and Flex-Cas12a samples was undetectable at predicted off-target sites (Supplementary Figure S5A, B), perhaps due in part to the use of RNP delivery instead of plasmids (45–47).

**Figure 5.**
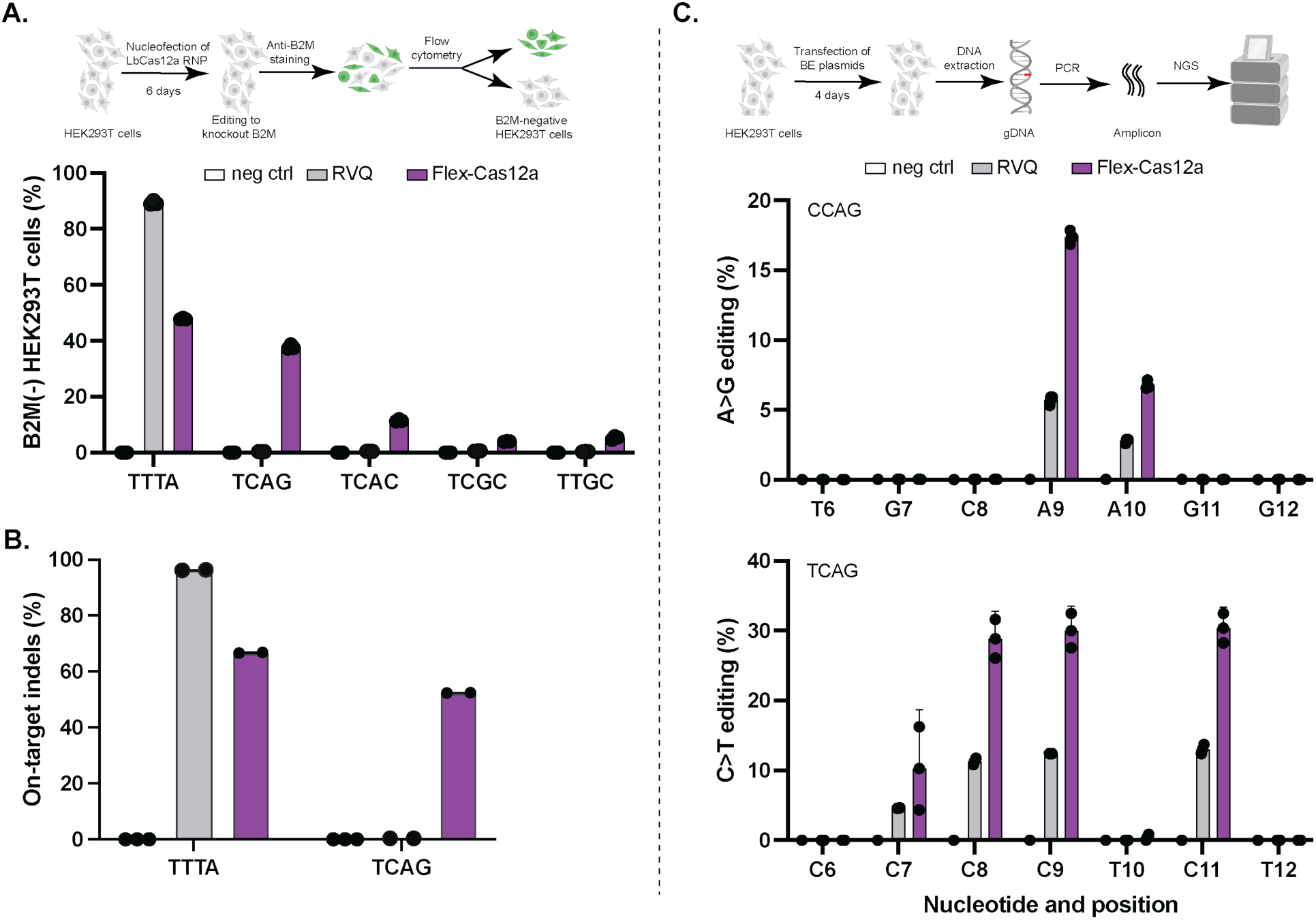
Genome editing at endogenous loci. **A.** Genome editing at endogenous loci in HEK293T cells. The upper panel illustrates the detection of genome editing in HEK293T cells using an anti-B2M antibody. The lower panel shows editing efficiencies at five endogenous loci in B2M gene. Flex-Cas12a can edit five endogenous loci with different PAMs, whereas LbCas12a-RVQ (RVQ) is restricted to the site with a canonical-PAM. TTTA is a canonical, while others, non-canonical PAMs. PAM sequence of each target is listed on X-axis. **B**. Genome editing analysis by next-generation sequencing (NGS). Editing efficiencies at two selected targets with a canonical PAM of 5’-TTTA-3’ and a non-canonical PAM of 5-TCAG-3’ are shown. Data presented here are generated from three independent technical replicates. Indels here represents insertions and deletions. **C**. Base editing analysis by NGS analysis. Top panel is a schematic illustration of analysis of base editing. Middle panel presents a representative data from adenine base editing (ABE). Bottom panel shows a representative data from cytosine base editing (CBE). All the data are presented as mean ± SD from three independent technical replicates.

Finally, we tested whether Flex-Cas12a could be employed for base editing (BE) at target sites with non-canonical PAMs. We fused either adenine base editor (ABE) or cytosine base editor (CBE) catalytic domains to either LbCas12a-RVQ or Flex-Cas12a to generate four base editor constructs: LbCas12a-RVQ-ABE, LbCas12a-RVQ-CBE, Flex-Cas12a-ABE and Flex-Cas12a-CBE. We also designed crRNAs to enable targeting of four endogenous genomic loci for ABE and for CBE, respectively. Following RNP delivery of these Cas12a base editors into HEK293T cells, NGS data showed that Flex-Cas12a-based base editors produced 2-3-fold higher editing efficiencies at sites with non-canonical PAMs compared to those generated by their LbCas12a-RVQ counterparts (Figure 5C). The four base-editor Flex-Cas12a constructs performed similarly to their wild-type counterparts at sites containing a canonical 5’-TTTV-3’ PAM (Figure 5C; Supplementary Figure S6A, B). Consistent with our other data, these results show that Flex-Cas12a not only matches activities of LbCas12a-RVQ at sites with a canonical PAM but can also edit efficiently at sites with non-canonical PAMs, expanding genome editing opportunities in mammalian cells. The low residual BE activity displayed by wild-type LbCas12a at non-canonical PAM sites observed in this study and others (21) may result from rapid deamination on dsDNA even without full *R*-loop formation (35, 48, 49). Taken together, these findings establish Flex-Cas12a as a versatile genome editing tool that enables modification of previously unreachable genomic loci, providing a robust new platform for both research and therapeutic applications.

## DISCUSSION

In this study, we used an unbiased directed evolution approach to identify seven PAM-relaxed LbCas12a variants with expanded DNA target recognition and genome editing capabilities. Among these, one derivative, Flex-Cas12a, was identified as the most effective as it displayed robust performance in both biochemical and cell-based assays. Flex-Cas12a recognizes a wide spectrum of 5’-NYHV-3’ PAMs, retains robust activity with the canonical 5’-TTTV-3’ PAM and has minimal off-target DNA binding. This expands the targetable human genome from approximately 1% to over 25%, representing a substantial improvement over previously engineered LbCas12a and AsCas12a variants, the best of which are theoretically restricted to targeting only ∼6% of the genome (18, 20, 21). Therefore, Flex-Cas12a enriches the genome editing toolbox, enabling precise targeting of previously inaccessible loci and facilitating both single-gene and multiplexed editing for genomic DNA manipulations.

To assess the potential mechanism of Flex-Cas12a’s expanded PAM recognition, we mapped all six mutations in this protein onto the crystal structure of the LbCas12a-crRNA-DNA ternary complex (PDB: 5XUS). We hypothesize that the D535G substitution alters interactions between the protein and DNA backbone in the PAM region (Figure 1B, bottom panel) and potentially broadens its PAM recognition. The introduced activity-enhancing mutations including G146R, R182V and E795Q (17) are located away from the PAM sequence and hence may primarily enhance enzymatic activity rather than altering PAM specificity, consistent with our results (Supplementary Figure S2C) (17). Taken together, Flex-Cas12a can target a wider array of PAMs without compromising catalytic activity.

Preliminary biochemical analysis (Figure 3A and 3B, Supplementary Figure S3C) also suggests that Flex-Cas12a avoids the off-target entrapment observed for variants of CRISPR-Cas9 with reduced PAM specificity (24, 26), enabling efficient target search and sustained specificity despite relaxed PAM constraints. Future biophysical analysis of these variants could reveal distinct energetic landscapes of target DNA recognition with the underpinning molecular determinants, enabling engineering of enzymes with higher versatility and efficiency. Coupled with LbCas12a’s genome editing utility (50, 51), Flex-Cas12a represents a significant addition to the genome editing toolbox.

## SUPPLEMENTARY DATA

Supplementary data are available at NAR Online.

## ACKNOWLEDGEMENTS

We acknowledge Ms. Netravathi Krishnappa (NGS Core Operations Manager and Sequencing Specialist, Center for Translational Genomics, Innovative Genomics Institute, UC Berkeley).

## Supplementary Figures

**Supplementary Figure S1.**
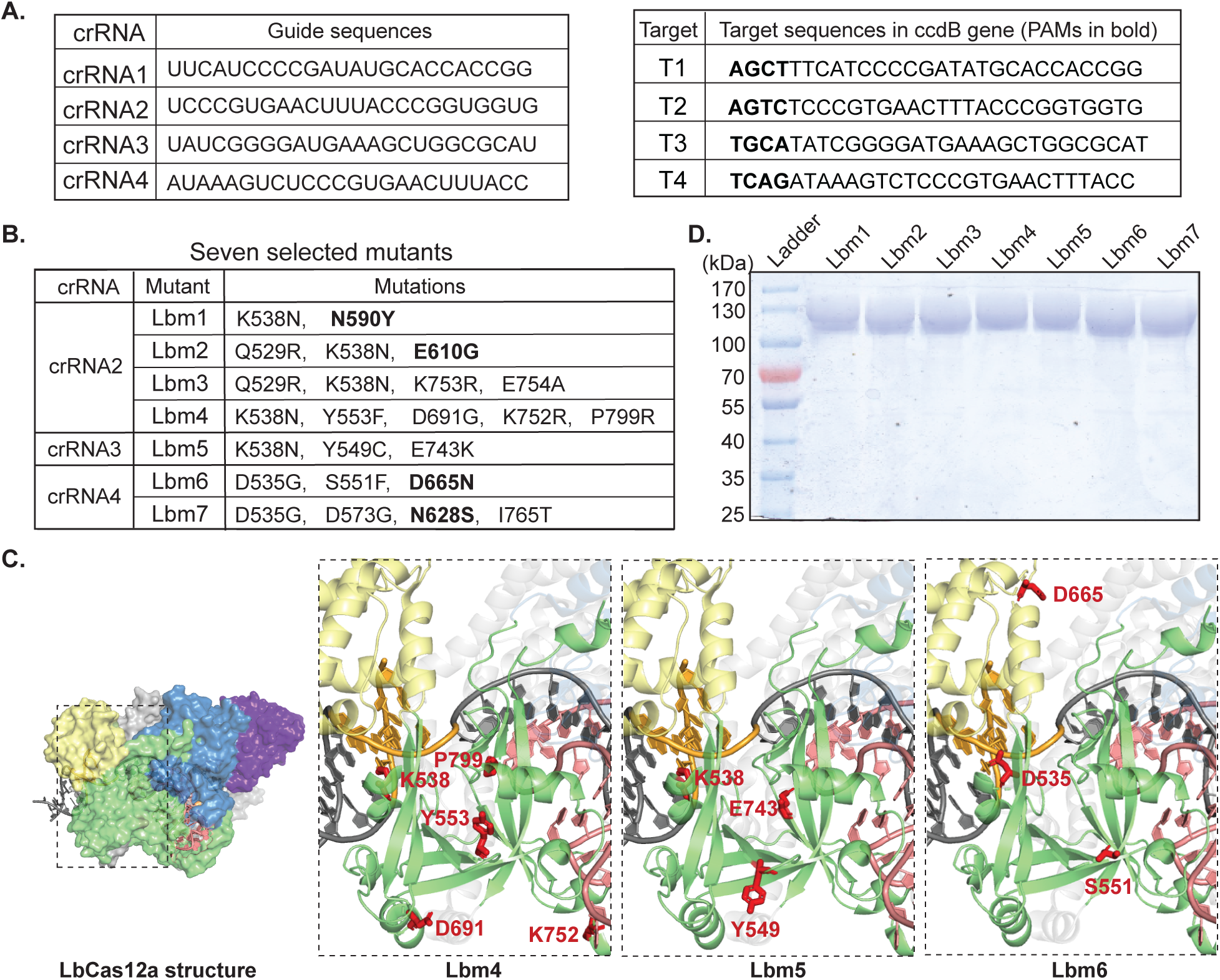
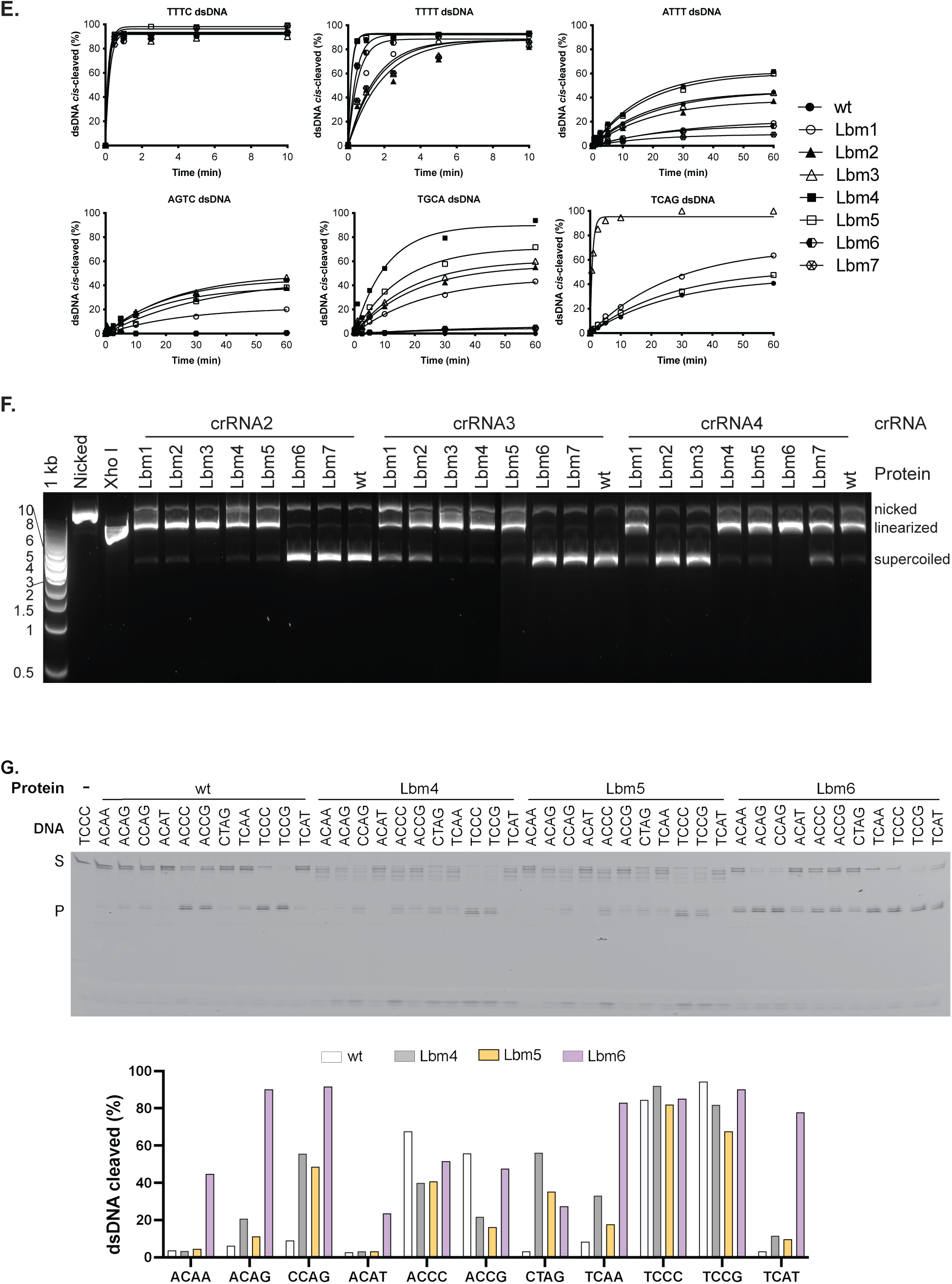
Generation and analysis of PAM-relaxed variants. **A.** Target sequences for directed evolution. Left panel shows the target sequences of four crRNAs (crRNA1 to 4) and right panel shows four *ccdB* target DNA sequences (presented as non-target strands) with randomly selected non-canonical PAMs (AGCT, AGTC, TGCA, or TCAG). **B.** Mutations in PAM-relaxed LbCas12a variants. Residues highlighted in bold indicate that they are localized within the PAM-interacting domain (PI). **C.** Structural presentation of the LbCas12a-dsDNA-crRNA ternary complex (PDB: 5XUS), highlighting mutation sites in Lbm4, Lbm5, and Lbm6. PI domain is colored in yellow, and WED domains are in green, PAM sequence is shown in brown, and all mutated residues are labeled in red. **D.** An SDS-PAGE gel image of the purified seven variants Lbm1-7. **E.** *In vitro* kinetic analysis of *cis*-cleavage activity. Cleavage efficiencies of wild-type (wt) and seven variants (Lbm1 to Lbm7) are shown. Data points represent the mean of two independent experiments. Target DNAs (derivatives of T0, listed in Table S1) used in these assays are same except PAM sequences. TTTC is a canonical PAM, while others, non-canonical PAMs. Each PAM sequence of DNA substrate is given on top of each panel. **F.** Plasmid cleavage assays of *ccdB*-containing plasmid DNA. Each designed crRNA targets a DNA sequence with a non-canonical PAM (crRNA2:AGTC, crRNA3:TGCA and crRNA4:TCAG). Each crRNA was tested against the seven Lbm1-7 variants as well as wild-type LbCas12a (wt). **G.** *In vitro* cleavage assays with various synthetic DNA substrates which are derived from DNA T0 with different non-canonical PAM sequences. The corresponding PAM sequences are listed on top of the gel image or on X-axis. Top panel presents a representative gel image. Lower panel quantifies cleavage efficiencies across eleven target DNAs by wt, Lbm4, Lbm5 and Lbm6, respectively. S presents substrate, and P, cleavage products.

**Supplementary Figure S2.**
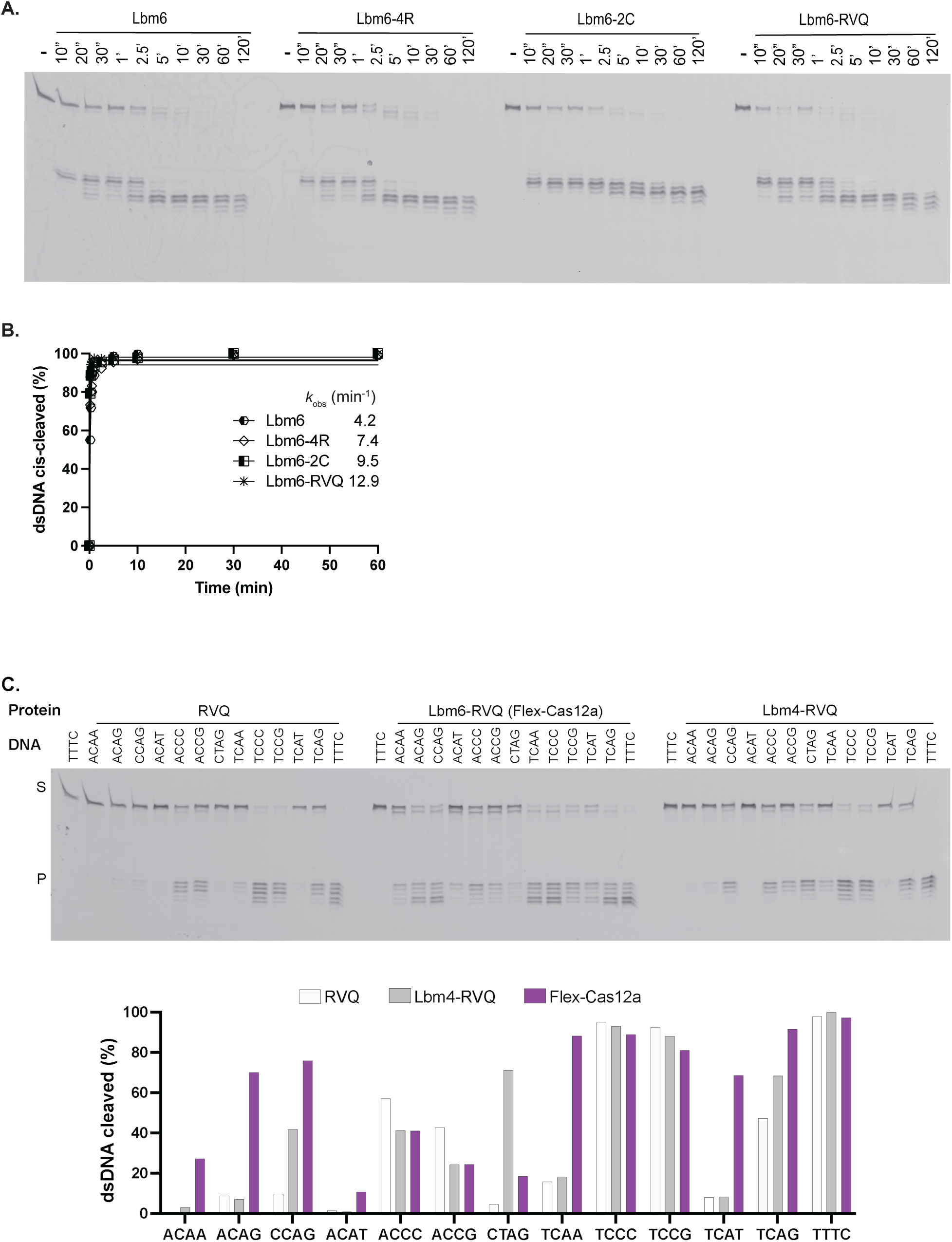
DNA cleavage assays by Lbm6 and its derivatives. **A.** A gel image of *in vitro* cleavages by Lbm6 and its derivatives of Lbm6-4R, Lbm6-2C and Lbm6-RVQ. Target DNA used in this assay is DNA T0 (listed in Table S1) with a PAM of 5’-TTTC-3’). **B.** Quantification of cleavage efficiencies of each variant over time. The cleavage results indicate that Lbm6-RVQ exhibits the highest activity and is renamed Flex-Cas12a from now on. **C.** *In vitro* cleavage assays of DNA substrates bearing different PAMs. Lbm4-RVQ, Flex-Cas12a and LbCas12a-RVQ (RVQ) were assessed. Upper panel shows a gel image of DNA cleavage reactions. Lower panel quantifies cleavage efficiencies for thirteen target DNAs with each protein. Target DNAs used in these assays are DNA T0 with different PAMs which are listed on top of the gel image or on X-axis. TTTC is a canonical PAM, while others, non-canonical PAMs. S presents substrate, and P, cleavage products.

**Supplementary Figure S3.**
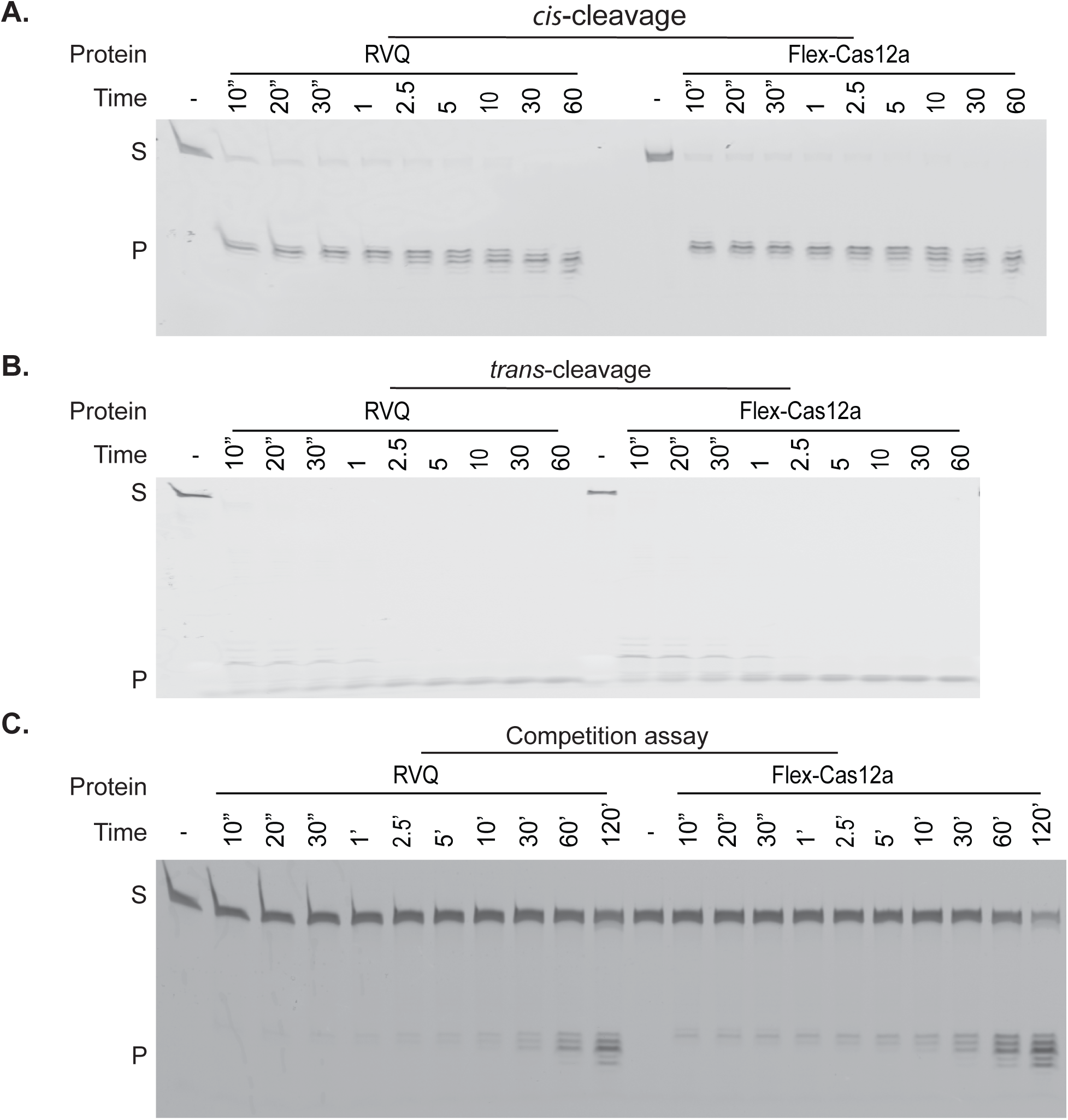
Comparison of DNA cleavage activities of Flex-Cas12a to LbCas12a-RVQ (RVQ). **A.** The gel image of *cis*-cleavage assays. DNA substrate used in assay is DNA T0 with a canonical PAM of 5’-TTTC-3’. **B.** The gel image of *trans*-cleavage assays. In this assay, 45 nM unlabeled target dsDNA T0 with a canonical PAM of 5’-TTTC-3’ was incubated with LbCas12a RNPs for 30 min at 37°C before addition of a labeled random ssDNA (no homology with the target DNAs or crRNAs, listed in Table S1). **C.** The gel image from the cleavage assay with a competitor DNA. In this assay with a competitor, 60 nM RNP, 10 nM labeled target DNA and 360 nM competitor DNA (pUC19 plasmid) were used. In this assay, labeled target DNA is DNA T0 with a canonical PAM of 5’-TTTC-3’.

**Supplementary Figure S4.**
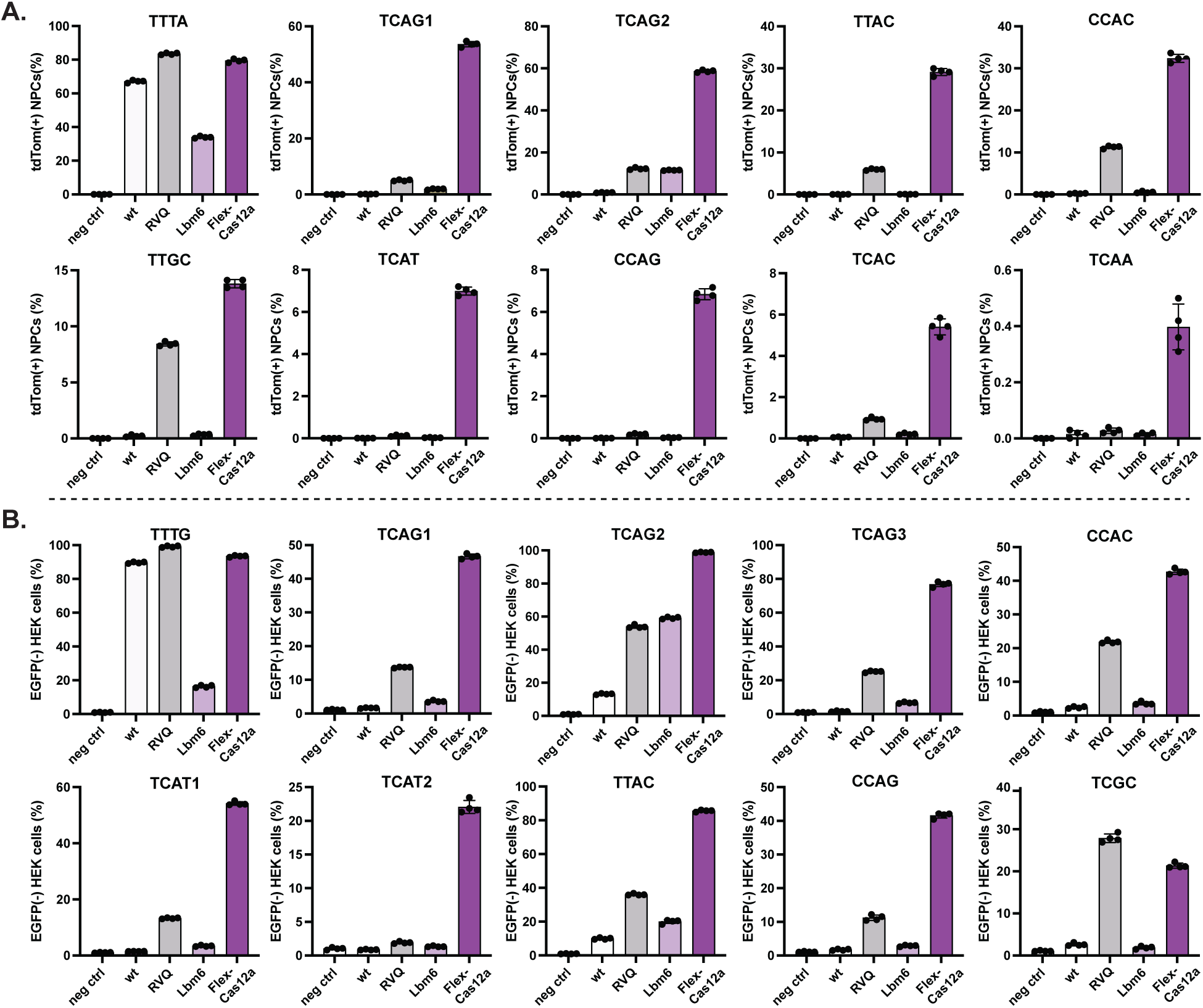
Genome editing in Ai9 tdTomato NPCs and HEK293-EGFP cells. **A.** Quantification of gnome editing at ten target sites in tdTomato NPCs. **B.** Quantification of genome editing at ten target sites in HEK293T-EGFP cells. PAM sequence for each target is listed on top of corresponding panel. In **A** and **B**, TTTA and TTTG are canonical PAMs, while others, non-canonical PAMs. All the editing data were quantified using flow cytometry and are presented as mean ± SD from four independent technical replicates. neg ctrl means the cells not treated with any proteins. Here, wt is abbreviated from wild-type, and RVQ, LbCas12a-RVQ.

**Supplementary Figure S5.**
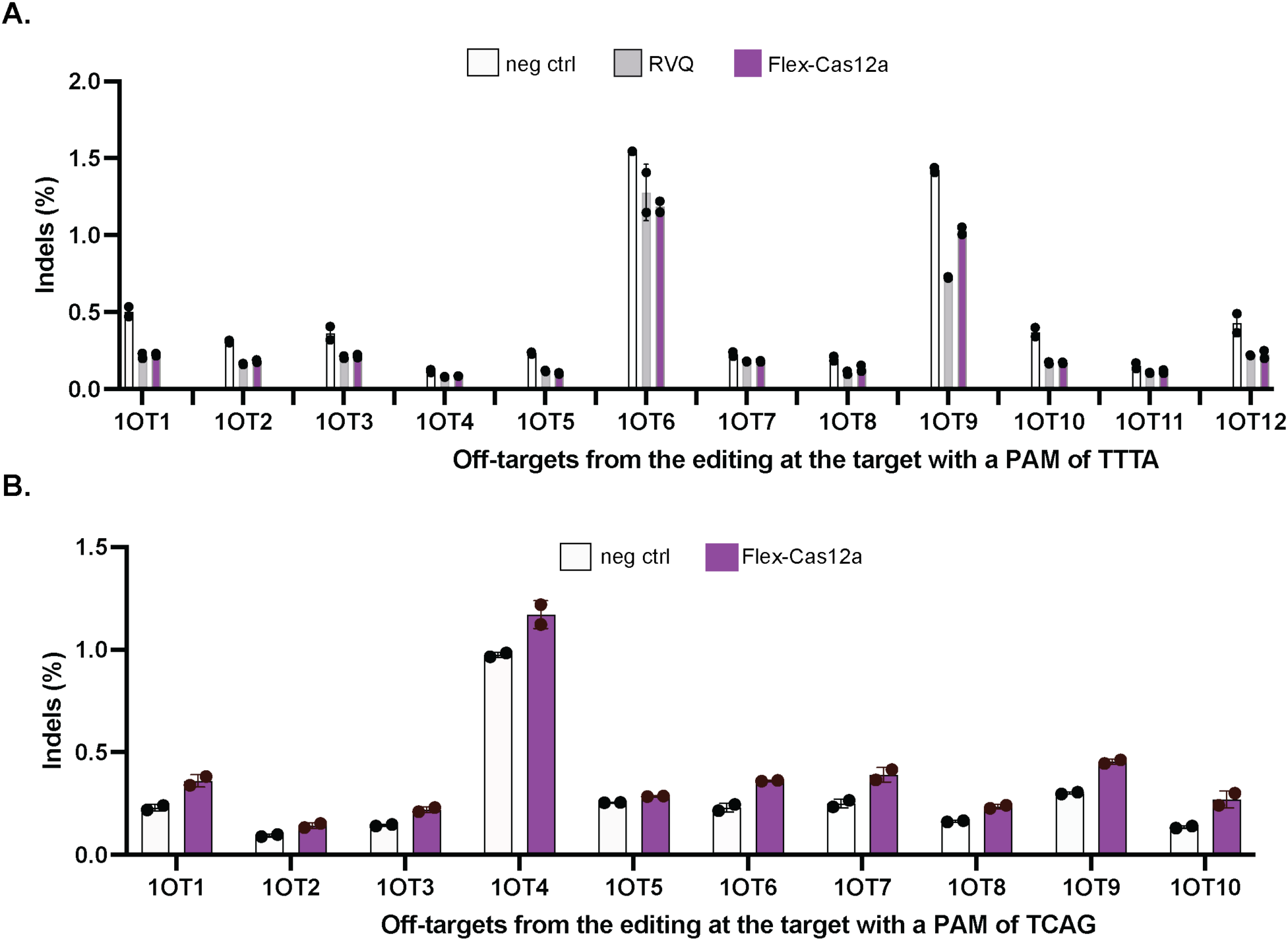
Off-target analysis of Flex-Cas12a. **A.** Off-target analysis of genomic DNAs isolated from the cells transfected with RNPs of LbCas12a-RVQ (RVQ) or Flex-Cas12a targeting a genomic site flanked with a 5’-TTTA-3’ PAM. Twelve off-target (OT) sites were analyzed. **B.** Off-target analysis of genomic DNAs isolated from the cells transfected with RNP of Flex-Cas12a targeting a genomic site flanked with a 5’-TCAG-3’ PAM. Ten off-target (OT) sites were analyzed. Each data point represents the average of two independent replicates. No off-target activity was detected in these assays. Neg ctrl means genomic DNAs isolated from untreated cells.

**Supplementary Figure S6.**
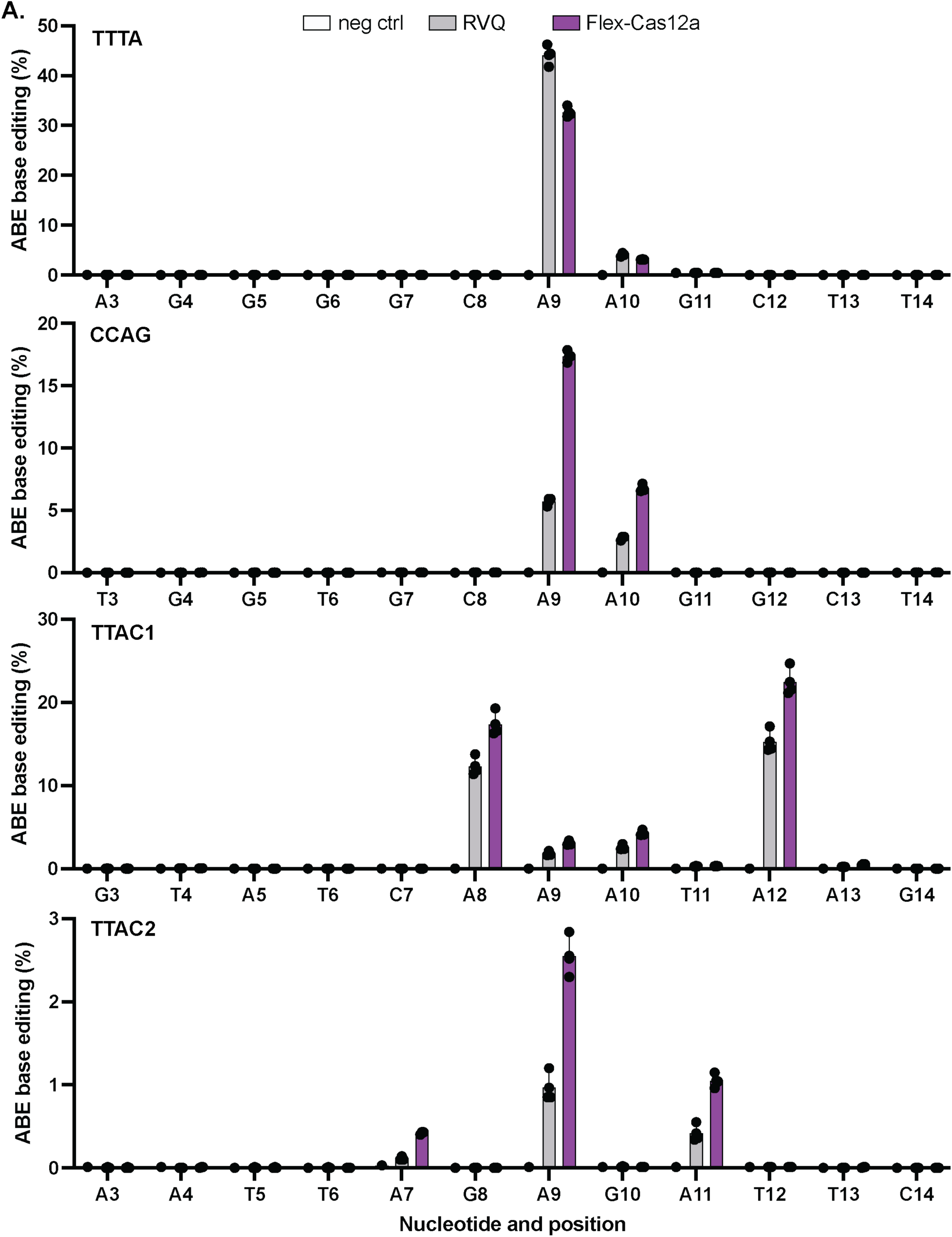
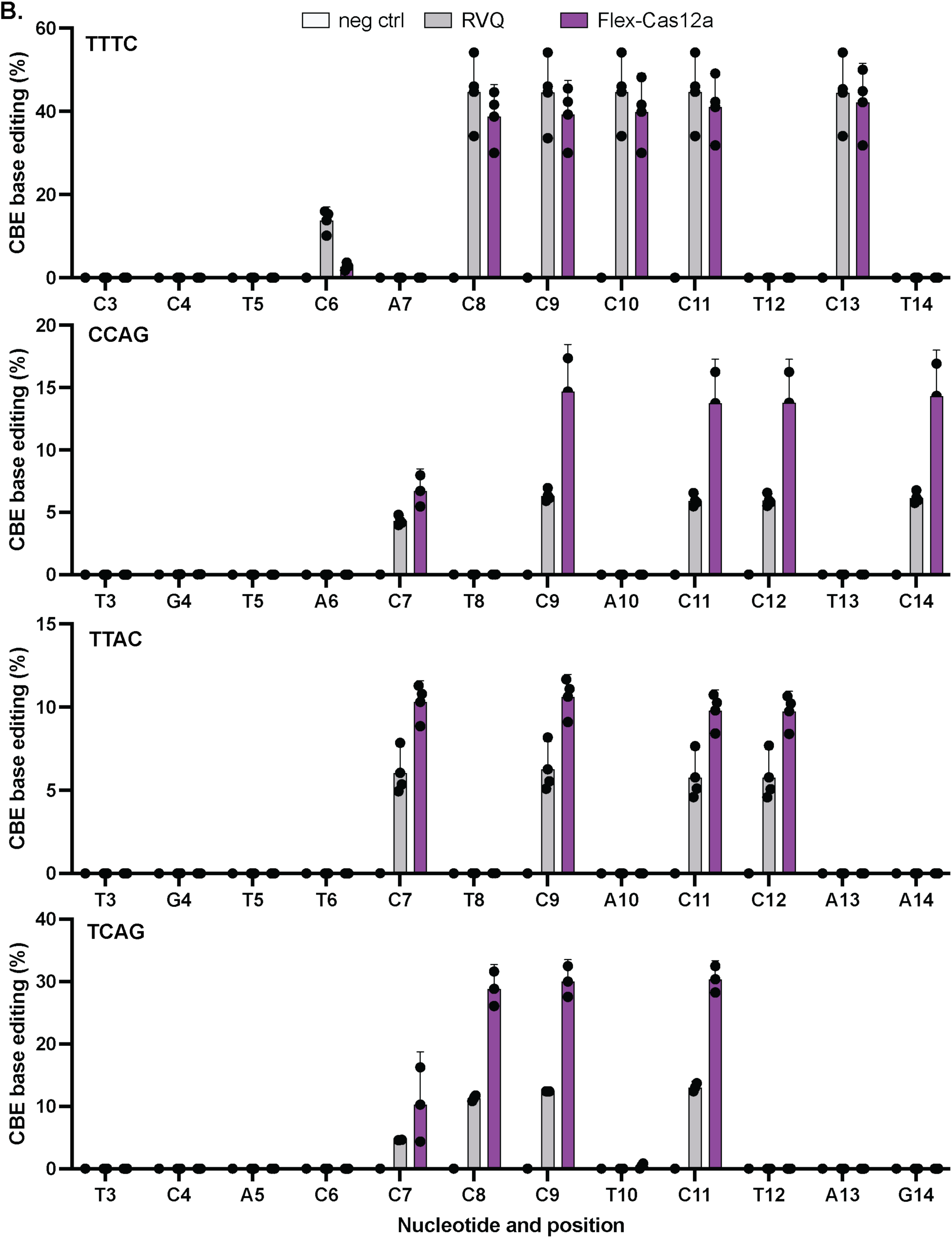
Base editing activity of Flex-Cas12a. **A.** Data from ABE base editing and **B.** Data from CBE base editing. Nucleotides at each position from 3 to 14 of target sequences are listed on X-axis. PAM sequence for each target is listed on top of each corresponding panel. TTTA and TTTC are canonical PAMs, while others, non-canonical PAMs. All the data are presented as mean ± SD from three independent replicates. RVQ is abbreviated from LbCas12a-RVQ.

