## Supplementary Figures for "Directed evolution expands CRISPR-Cas12a genome editing capacity"

**A.**

| crRNA | Guide sequences |
| --- | --- |
| crRNA1 | UUCAUCCCCGAUAUGCACCACCGG |
| crRNA2 | UCCCGUGAACUUUACCCGGUGGUG |
| crRNA3 | UAUCGGGGAUGAAAGCUGGCGCAU |
| crRNA4 | AUAAAGUCUCCCGUGAACUUUACC |

| Target | Target sequences in ccdB gene (PAMs in bold) |
| --- | --- |
| T1 | **AGCT**TTCATCCCCGATATGCACCACCGG |
| T2 | **AGTC**TCCCGTGAACTTTACCCGGTGGTG |
| T3 | **TGCA**TATCGGGGATGAAAGCTGGCGCAT |
| T4 | **TCAG**ATAAAGTCTCCCGTGAACTTTACC |

**B.**

Seven selected mutants

| crRNA | Mutant | Mutations |
| --- | --- | --- |
| crRNA2 | Lbm1 | K538N, **N590Y** |
| | Lbm2 | Q529R, K538N, **E610G** |
| | Lbm3 | Q529R, K538N, K753R, E754A |
| | Lbm4 | K538N, Y553F, D691G, K752R, P799R |
| crRNA3 | Lbm5 | K538N, Y549C, E743K |
| crRNA4 | Lbm6 | D535G, S551F, **D665N** |
| | Lbm7 | D535G, D573G, **N628S**, I765T |

**D.**

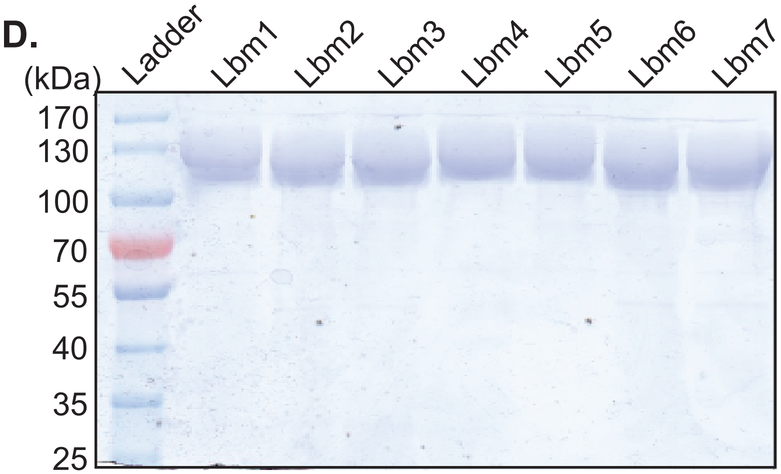


**C.**

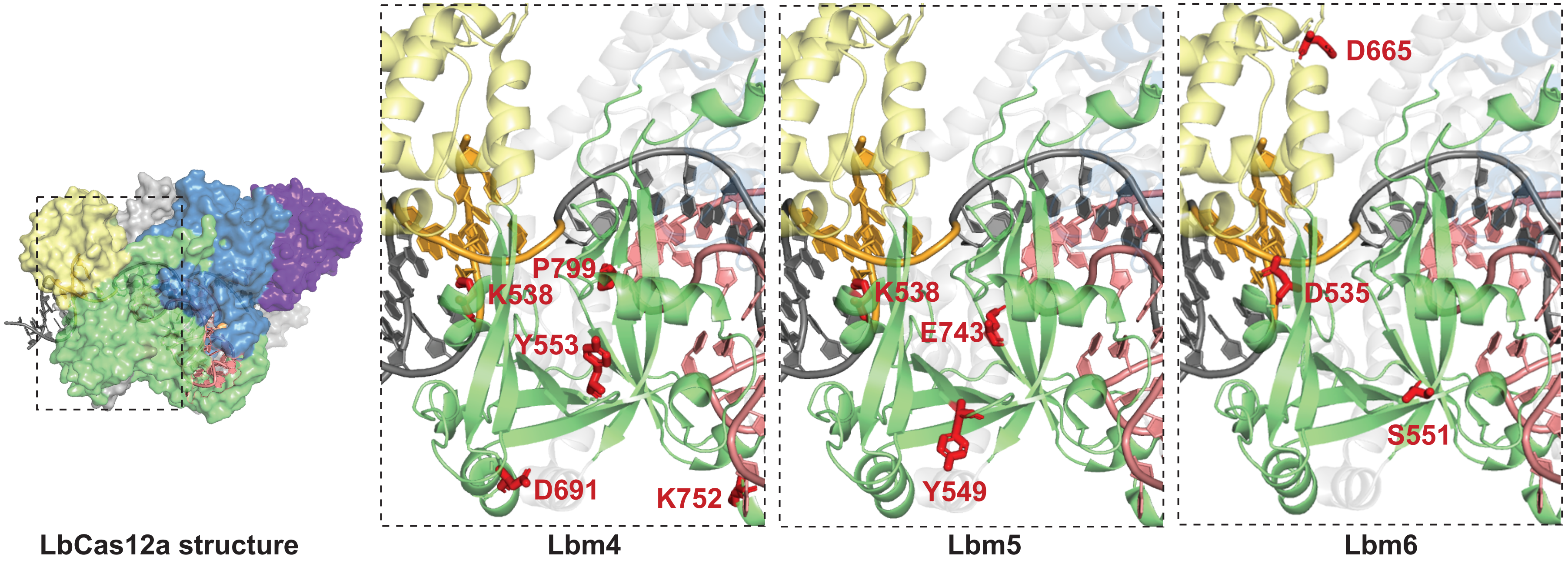


**Figure S1A - D**

E.

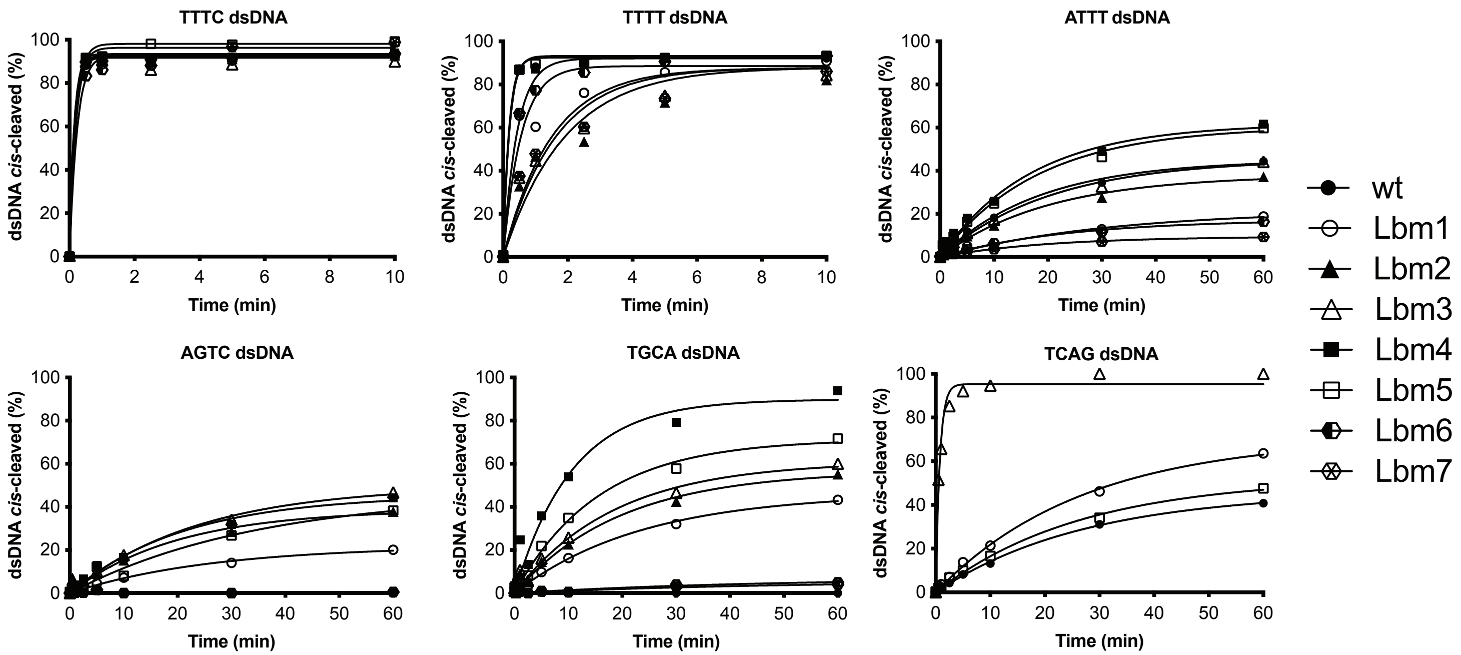


F.

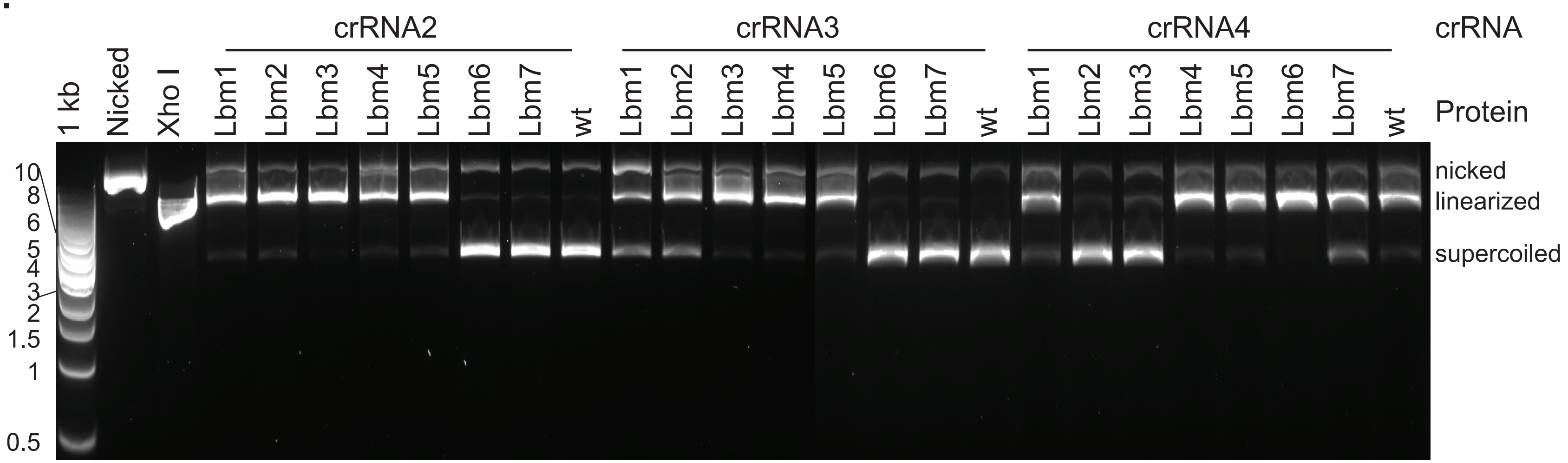


**Figure S1E - F**

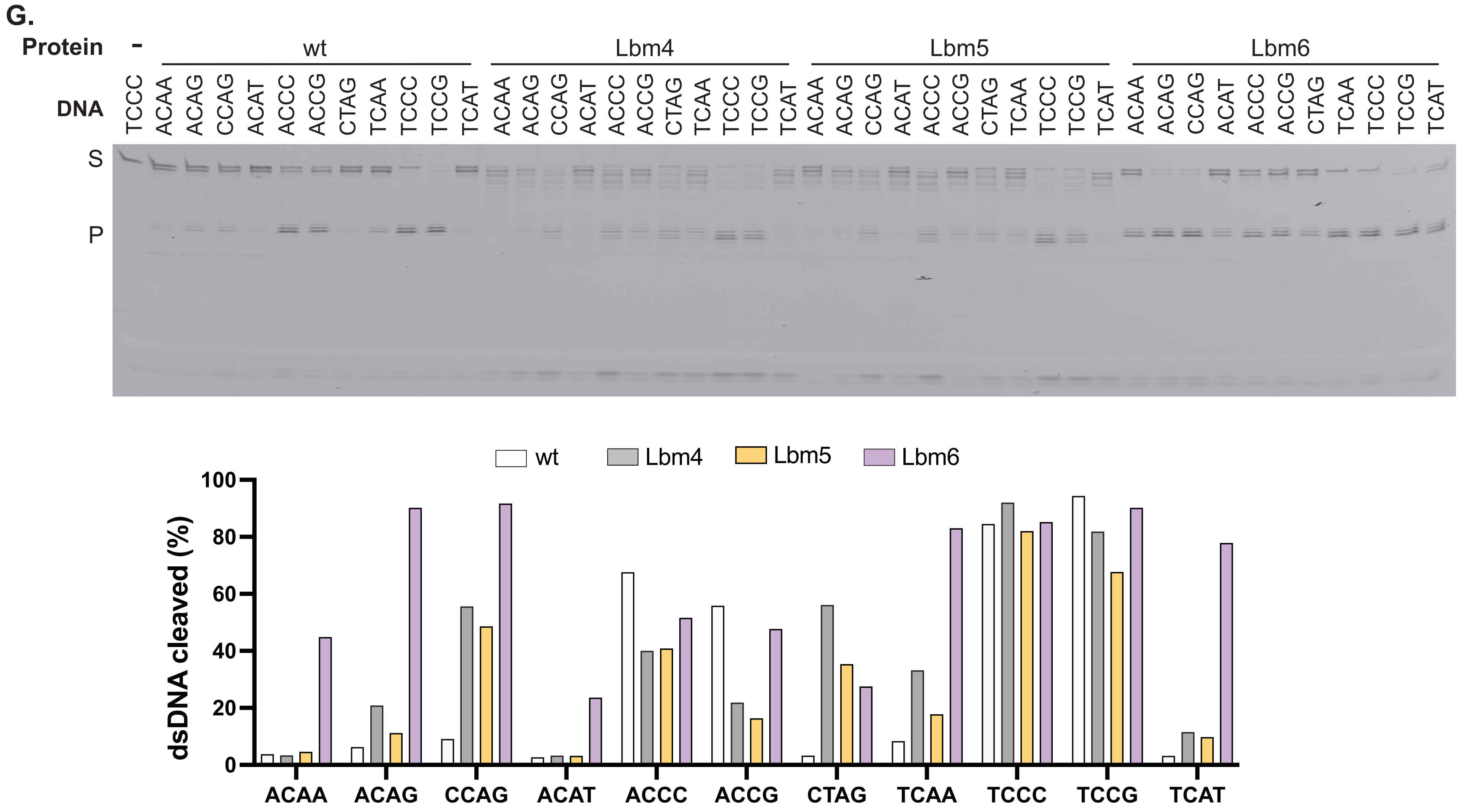
G.
Protein
-
wt
Lbm4
Lbm5
Lbm6
DNA
TCCC
ACAA
ACAG
CCAG
ACAT
ACCC
ACCG
CTAG
TCAA
TCCC
TCCG
TCAT
S
P
wt
Lbm4
Lbm5
Lbm6
dsDNA cleaved (%)
0
20
40
60
80
100
ACAA
ACAG
CCAG
ACAT
ACCC
ACCG
CTAG
TCAA
TCCC
TCCG
TCAT

**Figure S1G**

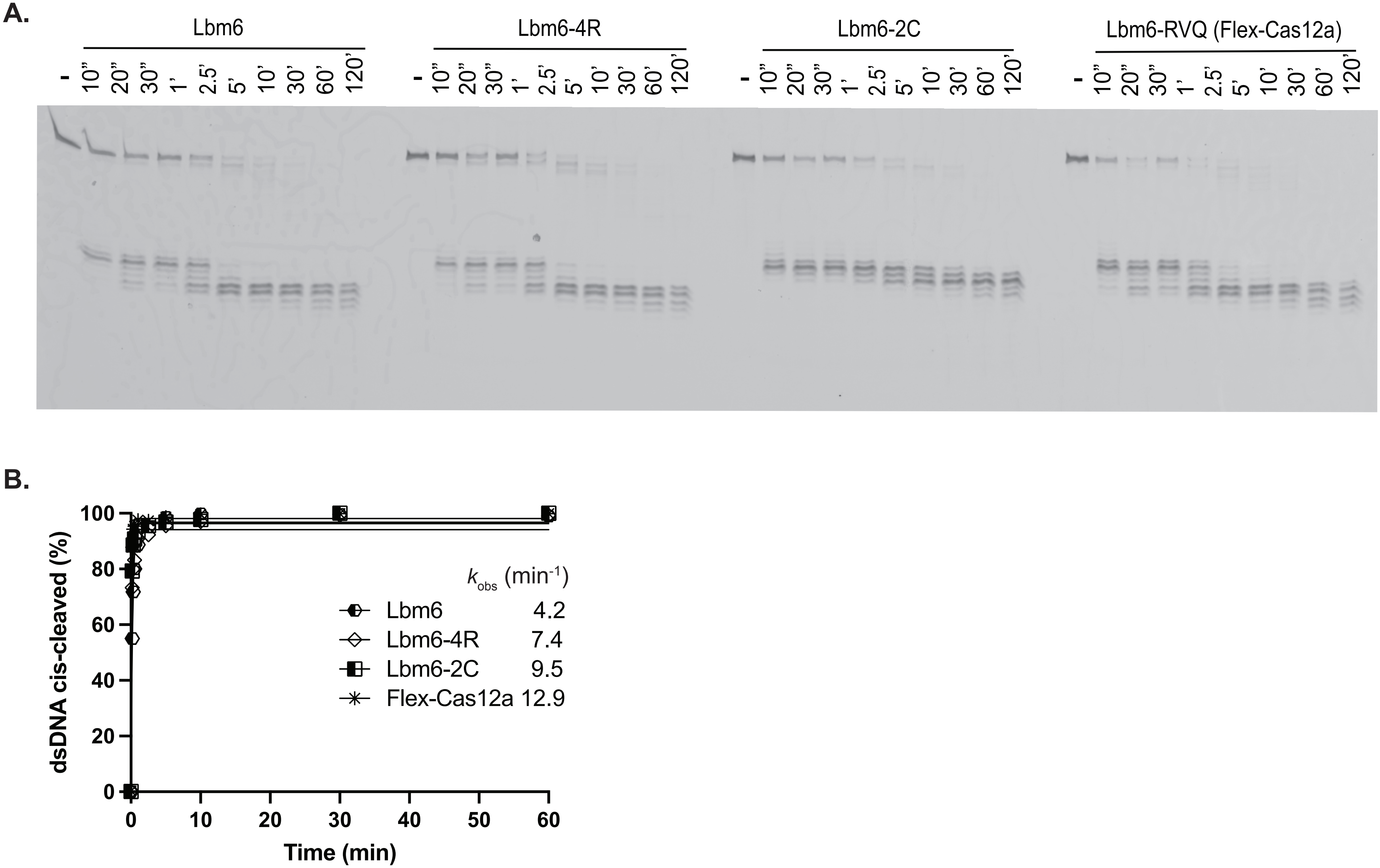


**Figure S2A and B**

C.

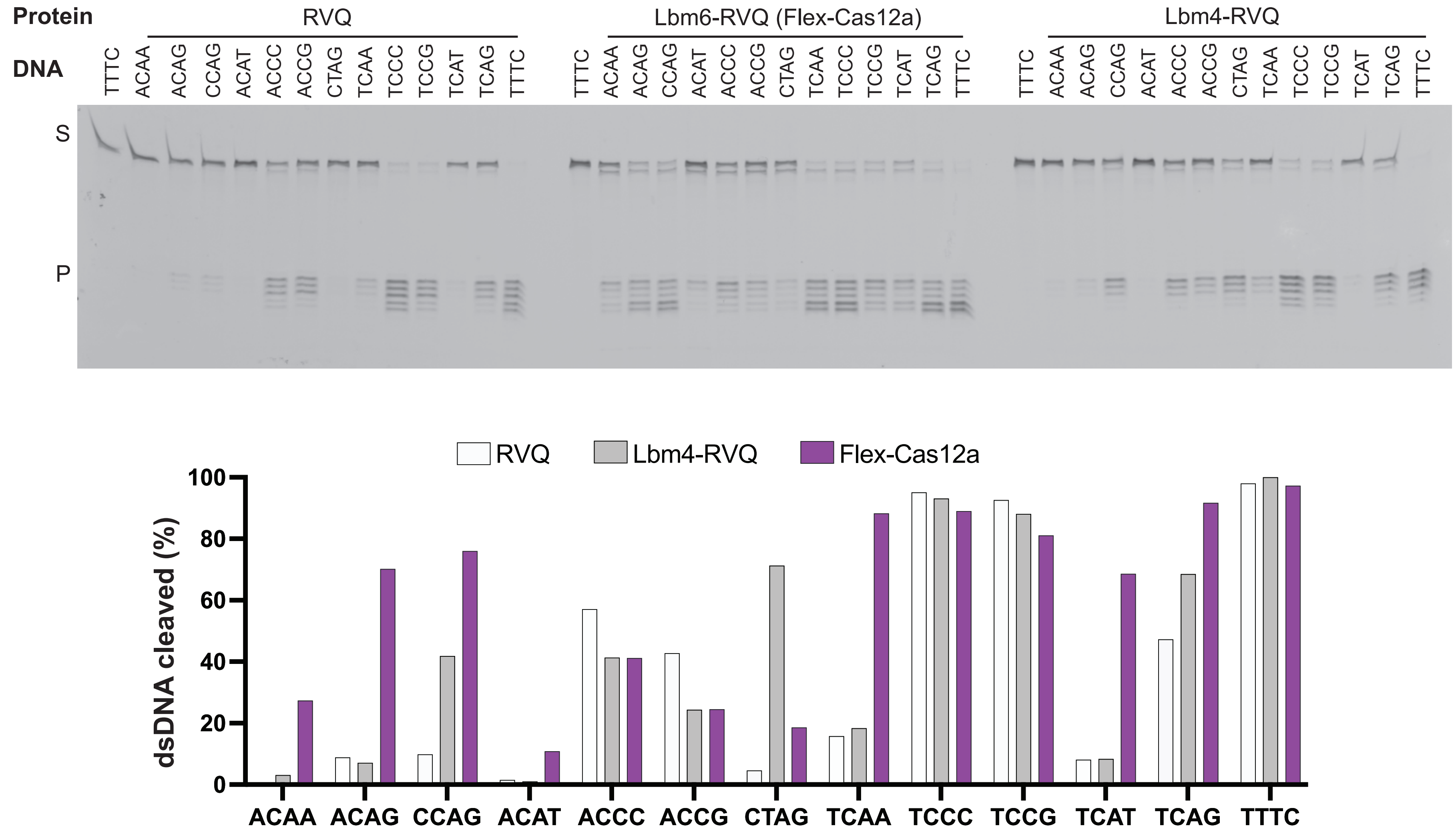
Protein
RVQ
Lbm6-RVQ (Flex-Cas12a)
Lbm4-RVQ
DNA
TTTC ACAA ACAG CCAG ACAT ACCC ACCG CTAG TCAA TCCC TCCG TCAT TCAG TTTC
S
P
RVQ
Lbm4-RVQ
Flex-Cas12a
dsDNA cleaved (%)
0
20
40
60
80
100
ACAA ACAG CCAG ACAT ACCC ACCG CTAG TCAA TCCC TCCG TCAT TCAG TTTC

**Figure S2C**

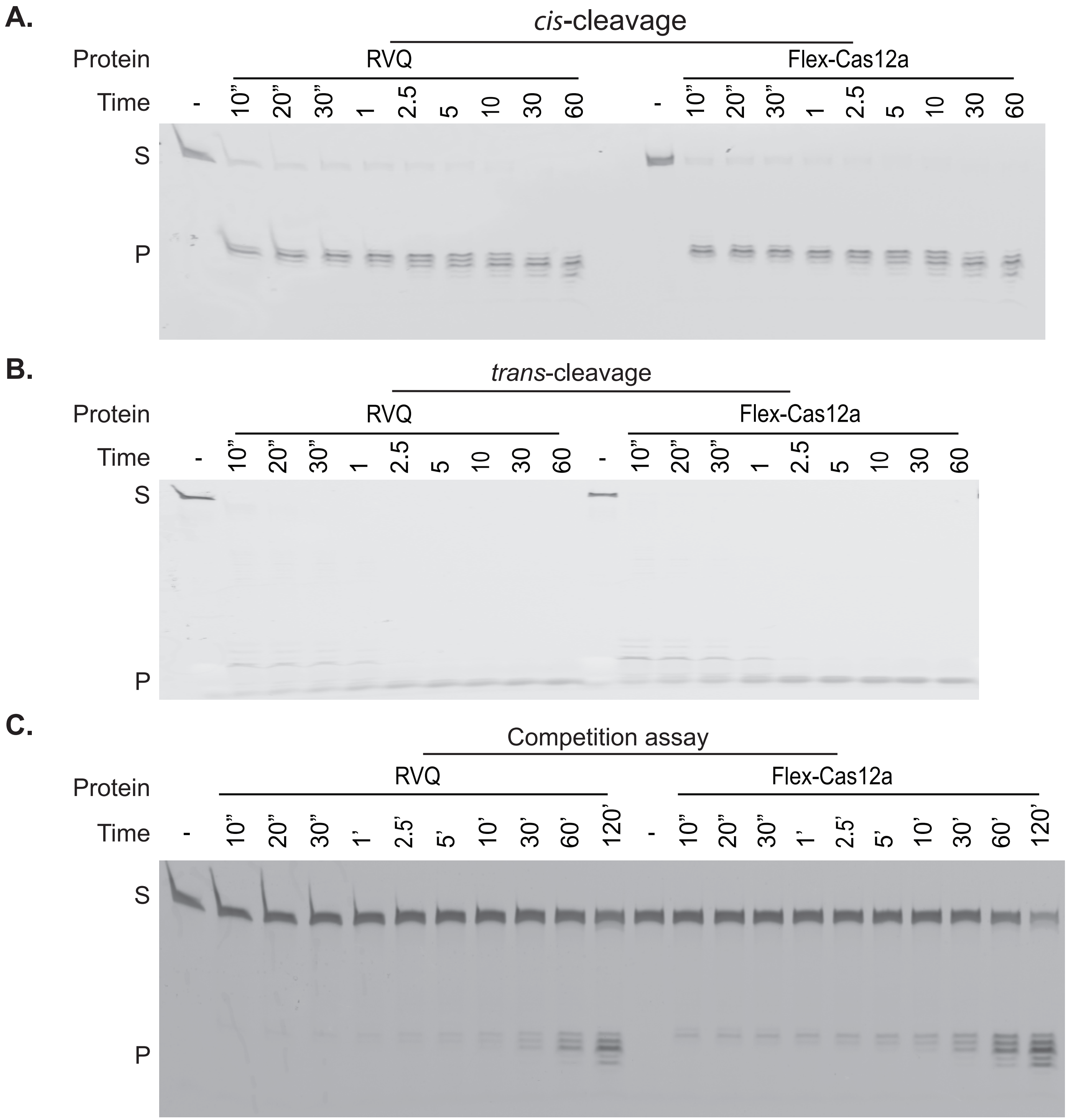
A.
*cis*-cleavage
Protein
RVQ
Flex-Cas12a
Time
-
10”
20”
30”
1
2.5
5
10
30
60
S
P
B.
*trans*-cleavage
C.
Competition assay
10’
30’
60’
120’
1’
2.5’
5’

**Figure S3**

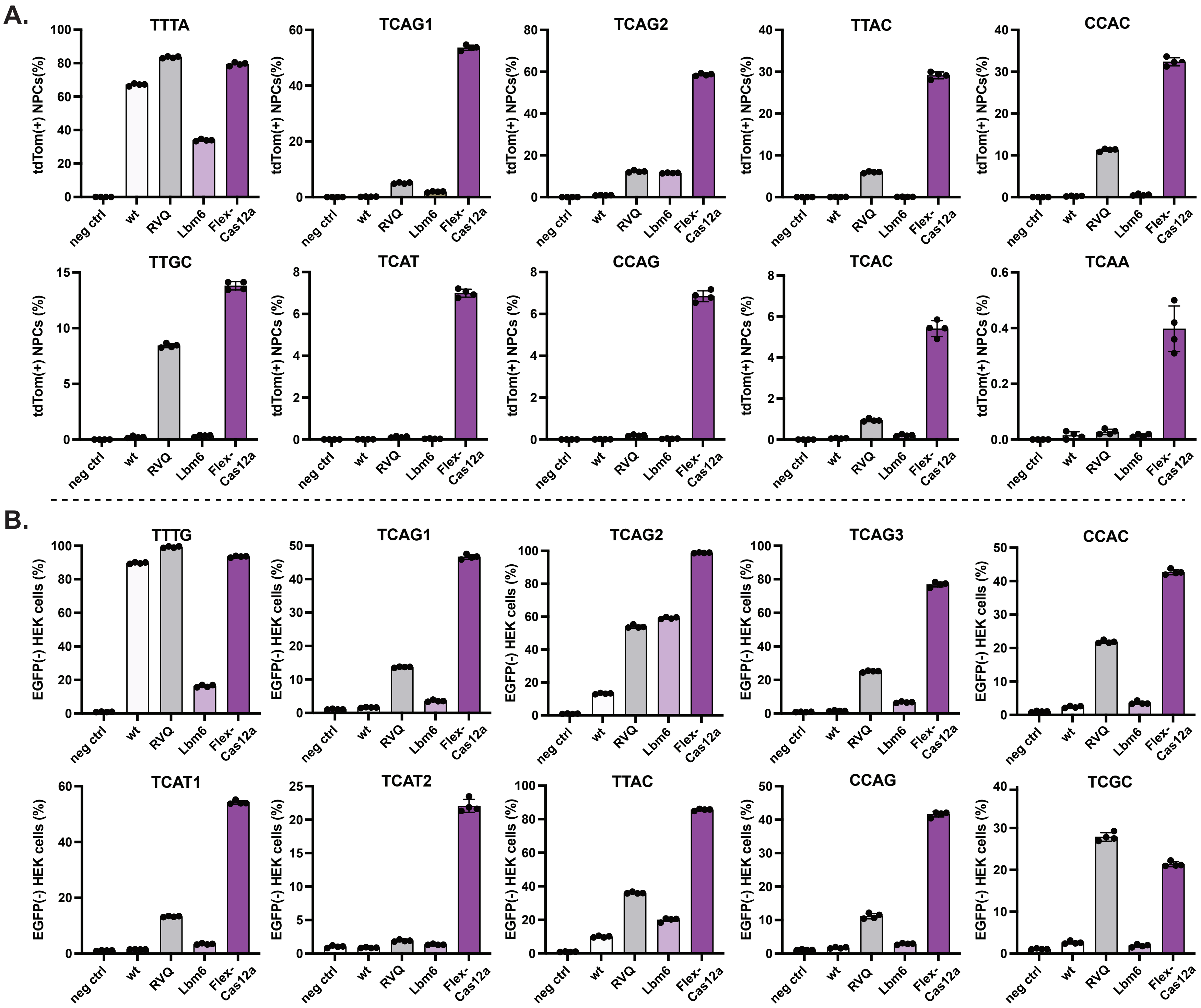
A.
TTTA
TCAG1
TCAG2
TTAC
CCAC
TTGC
TCAT
CCAG
TCAC
TCAA
tdTom(+) NPCs(%)
tdTom(+) NPCs (%)
B.
TTTG
TCAG1
TCAG2
TCAG3
CCAC
TCAT1
TCAT2
TTAC
CCAG
TCGC
EGFP(-) HEK cells (%)
neg ctrl
wt
RVQ
Lbm6
Flex-Cas12a

Figure S4

A.

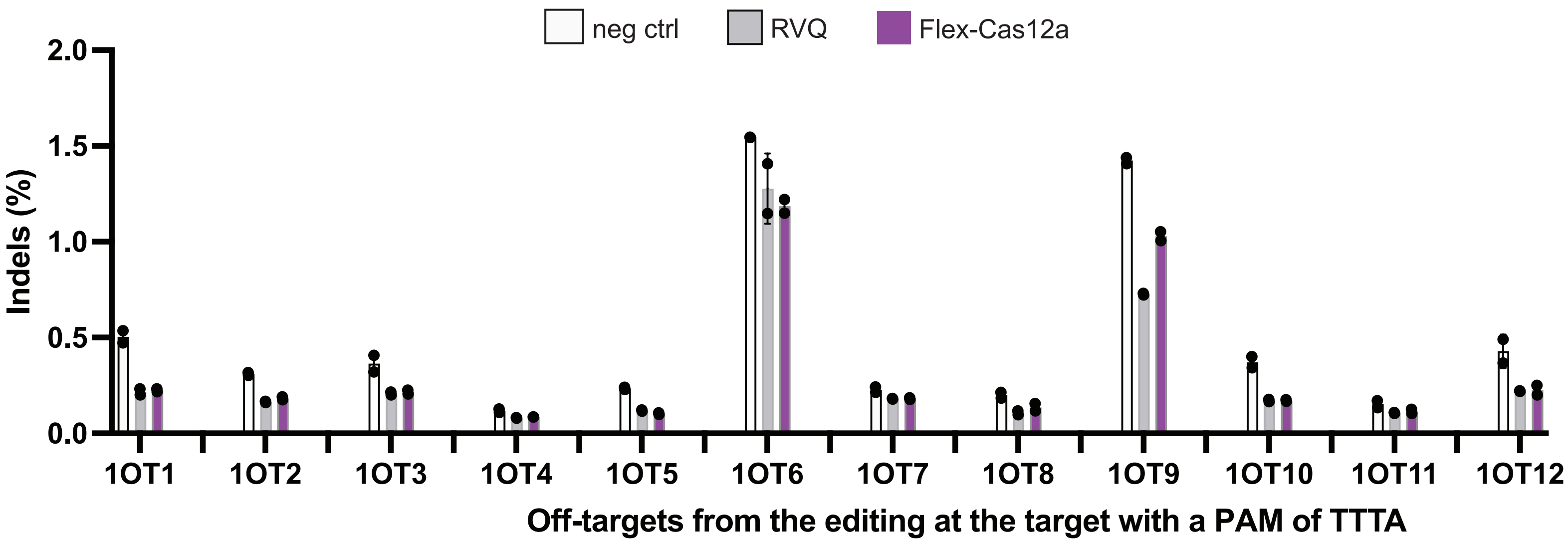


B.

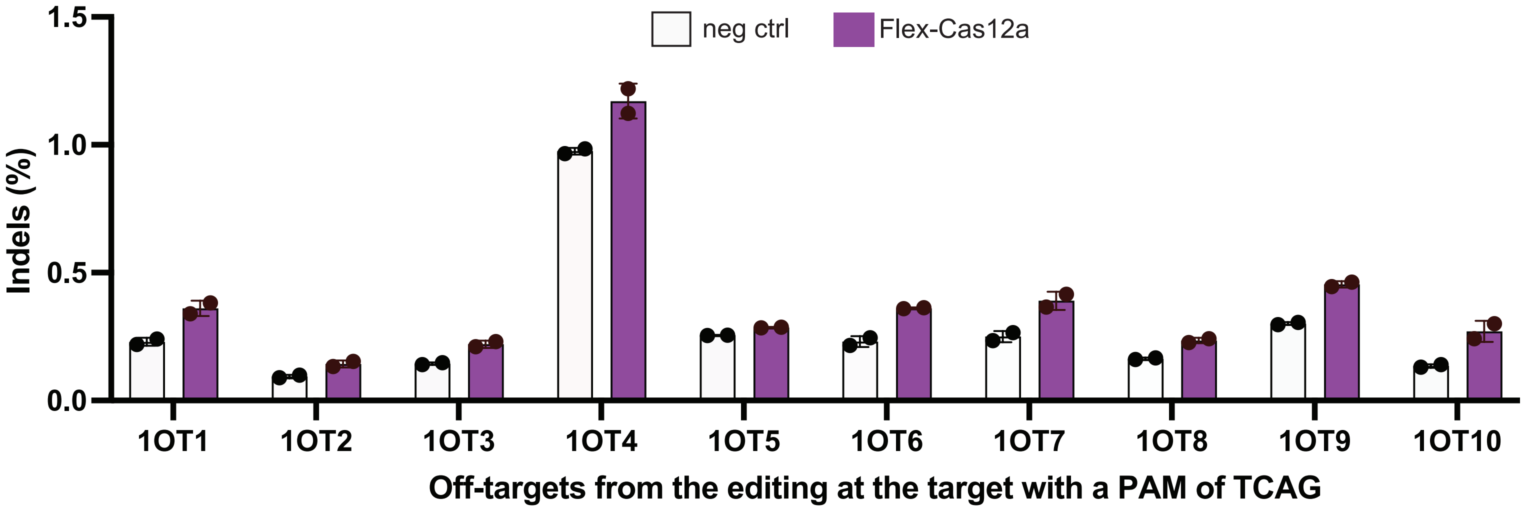


Figure S5A and B

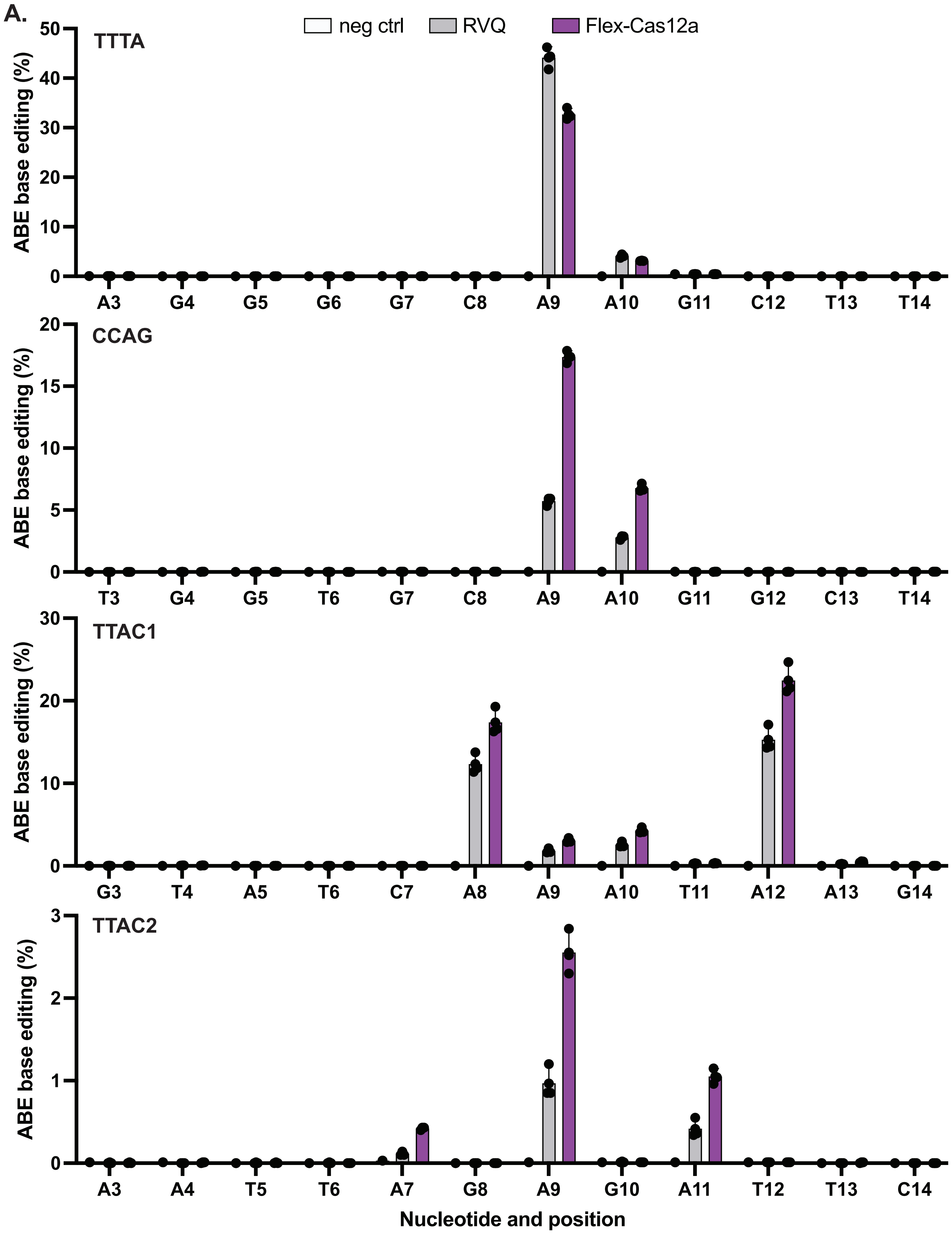

A.
neg ctrl
RVQ
Flex-Cas12a
TTTA
CCAG
TTAC1
TTAC2
ABE base editing (%)
0
10
20
30
40
50
0
5
10
15
20
0
10
20
30
0
1
2
3
A3 G4 G5 G6 G7 C8 A9 A10 G11 C12 T13 T14
T3 G4 G5 T6 G7 C8 A9 A10 G11 G12 C13 T14
G3 T4 A5 T6 C7 A8 A9 A10 T11 A12 A13 G14
A3 A4 T5 T6 A7 G8 A9 G10 A11 T12 T13 C14
Nucleotide and position


Figure S6A

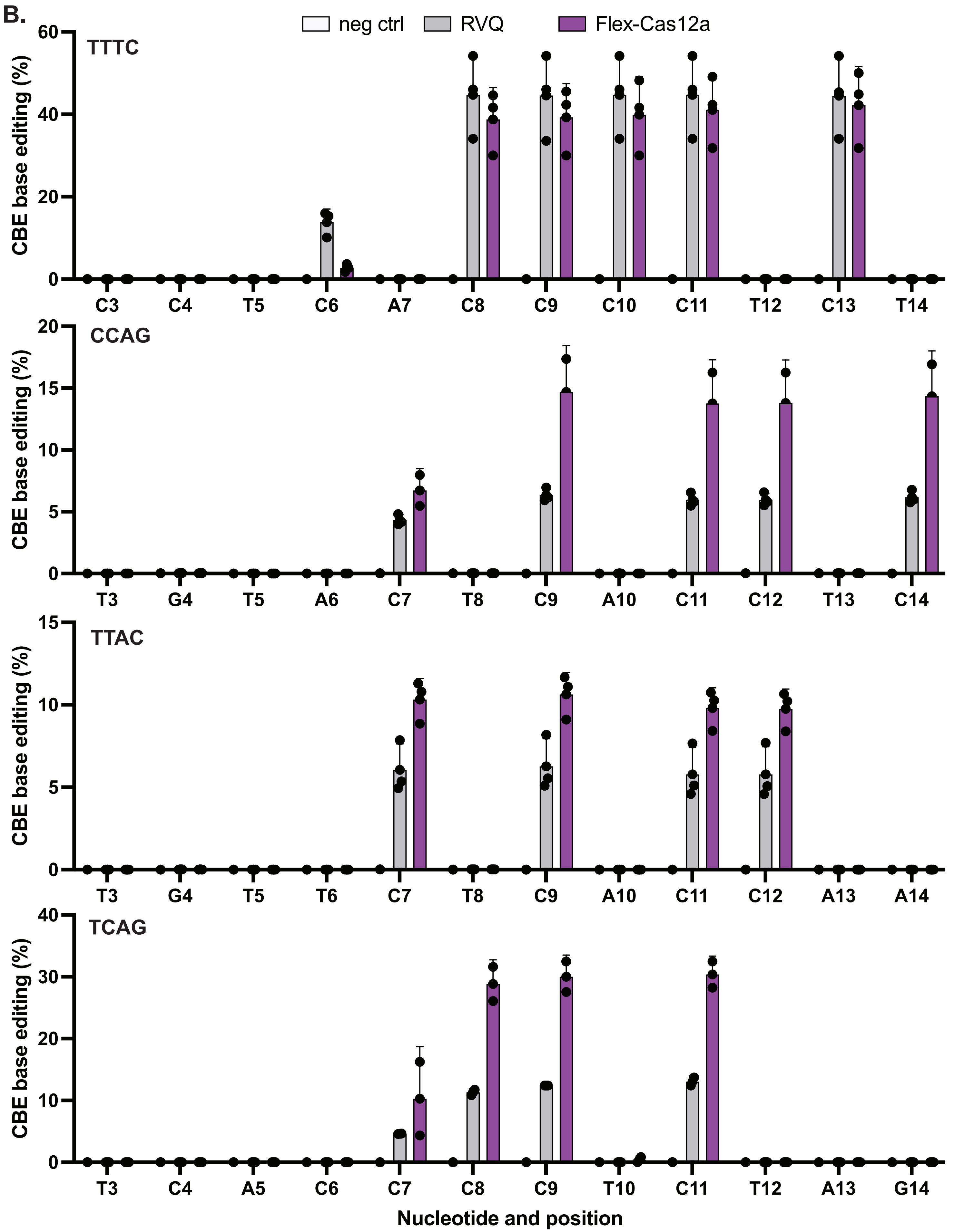
B.
neg ctrl
RVQ
Flex-Cas12a
TTTC
CCAG
TTAC
TCAG
CBE base editing (%)
C3 C4 T5 C6 A7 C8 C9 C10 C11 T12 C13 T14
T3 G4 T5 A6 C7 T8 C9 A10 C11 C12 T13 C14
T3 G4 T5 T6 C7 T8 C9 A10 C11 C12 A13 A14
T3 C4 A5 C6 C7 C8 C9 T10 C11 T12 A13 G14
Nucleotide and position

Figure S6B
